# Preclinical characterization and target validation of the antimalarial pantothenamide MMV693183

**DOI:** 10.1101/2021.05.12.443866

**Authors:** Laura E. de Vries, Patrick A.M. Jansen, Catalina Barcelo, Justin Munro, Julie M.J. Verhoef, Charisse Flerida A. Pasaje, Kelly Rubiano, Josefine Striepen, Judith M. Bolscher, Rob Henderson, Tonnie Huijs, Karin M.J. Koolen, Patrick K. Tumwebaze, Tomas Yeo, Anna C.C. Aguiar, Iñigo Angulo-Barturen, Alisje Churchyard, Jake Baum, Benigno Crespo Fernández, Francisco-Javier Gamo, Rafael V.C. Guido, María Belén Jiménez-Diaz, Dhelio B. Pereira, Rosemary Rochford, Laura M. Sanz, Graham Trevitt, Sergio Wittlin, Roland A. Cooper, Philip J. Rosenthal, Robert W. Sauerwein, Joost Schalkwijk, Pedro H.H. Hermkens, Roger Bonnert, Brice Campo, David A. Fidock, Manuel Llinás, Jacquin C. Niles, Taco W.A. Kooij, Koen J. Dechering

## Abstract

Drug resistance and a dire lack of transmission-blocking antimalarials hamper malaria elimination. Here, we present the pantothenamide MMV693183 as a first-in-class acetyl-CoA synthetase (ACS) inhibitor to enter preclinical development. Our studies demonstrated attractive drug-like properties and *in vivo* efficacy in a humanized mouse model of *Plasmodium falciparum* infection. The compound showed exceptional *in vitro* activity against *P. falciparum* and *P. vivax* clinical isolates, and potently blocked *P. falciparum* transmission to *Anopheles* mosquitoes. Genetic and biochemical studies identified ACS as the target of the MMV693183-derived antimetabolite, CoA-MMV693183. MMV693183 was well adsorbed after oral administration in mice, rats and dogs. Pharmacokinetic – pharmacodynamic modelling predicted that a single 30 mg oral dose is sufficient to cure a malaria infection in humans. In conclusion, the ACS-targeting compound MMV693183 represents a promising addition to the portfolio of antimalarials in (pre)clinical development with a novel mode of action for the treatment of malaria and blocking transmission.

## Introduction

Malaria remains a significant global infectious disease, caused by parasites of the genus *Plasmodium*. In the past two decades there was a major decline in malaria cases and deaths, however, this progress has slowed, indicating the need for new interventions (1). Drug resistance against many front-line therapies is emerging and spreading around the world, threatening the efficacy of these drugs (1–3). There is an urgent need for new therapeutics to combat the spread of resistance and progress towards malaria elimination. Target product profiles and target candidate profiles were developed to guide the discovery and clinical development of new antimalarials (4). Current approaches for new malaria treatments aim for a combination of two or more inexpensive, potent, fast-acting molecules that act on multiple parasite stages and provide a single dose cure (4). Compounds with new modes of action are favored, since no pre-existing resistance in the field would be expected.

Coenzyme A (CoA) is required for numerous processes within the cell, including lipid synthesis, protein acetylation and energy supply, and it is highly conserved among prokaryotes and eukaryotes (5). *Plasmodium* parasites rely on this pathway by uptake of the essential nutrient pantothenic acid (pantothenate or vitamin B5) (6, 7) that is converted into CoA in five enzymatic reactions (7). The CoA biosynthesis pathway in *Plasmodium* species has been considered a potential drug target since the discovery of the antimicrobial activity of pantothenic acid derivatives in the 1940s (8). Different libraries of pantothenic acid derivatives have been synthesized since then (8), however, due to poor stability in human serum they have never been developed into clinical candidates (9–11).

In the past decade, a focus has been on developing stable pantothenamides, in which the terminal carboxyl group of pantothenic acid is replaced by amides (11–13), including our recently synthesized pantothenamides with an inverted-amide bond (PanAms) (14). These PanAms are highly potent against pathology-causing asexual blood stages and transmittable gametocytes, consistent with the essentiality of several enzymes of the CoA pathway in both life-cycle stages (15–17). This indicates their potential to be developed into antimalarials that target a wholly novel pathway thereby curing the disease and block transmission to the mosquito host.

The exact mechanism of action of PanAms has been debated extensively, with pantothenate uptake, pantothenate kinase or CoA-utilizing processes as possible targets (7, 14, 17–19). The latest studies have indicated that the latter is the likely target. PanAms are metabolized, and form analogs of CoA pathway metabolites, including 4’P-PanAm, dP-CoA-PanAm and CoA-PanAm (14, 17, 18). A combination of biochemical and genetic approaches have demonstrated that these antimetabolites likely target the downstream enzymes acetyl-CoA synthetase (ACS; PF3D7_0627800) and acyl-CoA synthetase 11 (ACS11; PF3D7_1238800), thereby inhibiting the synthesis of acetyl-CoA (14). However, definitive proof of drug-enzyme interactions remained elusive.

Here, we describe the generation of the novel pantothenamide MMV693183 and demonstrate that its CoA-PanAm metabolite targets ACS. Moreover, MMV693183 has improved potency and metabolic stability, and a prolonged killing effect in a humanized mouse model of *P. falciparum* in comparison to previously described PanAms, and thus meets the requirements for further (pre)clinical development (4).

## Results

### MMV693183 is a potent antimalarial drug candidate

We recently synthesized a novel class of PanAms with an inverted-amide bond that resulted in compound MMV689258 with a limited predicted half-life in humans (14). Therefore, we continued chemical optimization, and a subseries of potent compounds with the aryl directly coupled to the inverted amide emerged (Table 1) (20). These PanAms showed activity against asexual and sexual blood-stage *P. falciparum* parasites with IC_50_ values ≤9.4 nM and ≤31 nM, respectively, except for one PanAm that was not active up to 1 μM against gametocytes (Table 1). To test *in vivo* activity, humanized mice were infected with *P. falciparum* and treated with a single dose of 50 mg/kg of PanAm by oral gavage. For all compounds, blood concentrations decreased rapidly over time and were either below or near the detection limit (5 ng/ml) after 24 h. In spite of the rapid elimination, MMV1542001, MMV884962, and MMV693183 reduced parasitemias below detectable levels over the course of three days, while parasitemias were not fully cleared upon treatment with MMV689258, MMV693182, and MMV976394 (Fig 1a; Table 1; Table S1). All five new PanAms had improved metabolic stability in a human primary hepatocyte relay assay compared to MMV689258 (Table 1) (21), potentially leading to a longer half-life. Attempts to obtain crystalline forms of the six PanAms, important for future drug formulation in tablets, were only successful for MMV689258, MMV693183, and MMV693182 (Fig S1), and resulted in favorable melting temperatures for the latter two compared to MMV689258 (Table 1). MMV693183 was selected as an advanced lead compound for further study, as it combined all improved characteristics. Furthermore, it was highly soluble in PBS, as well as in fasted- and fed-state simulated intestinal fluids (7.1, 9.2, 9.1 mg/ml, respectively) (Table S2). MMV693183 was also chemically stable after storage under stress conditions (40 °C, 75% relative humidity in an open and closed container or at 60 °C) (Fig S2).

**Fig 1.**
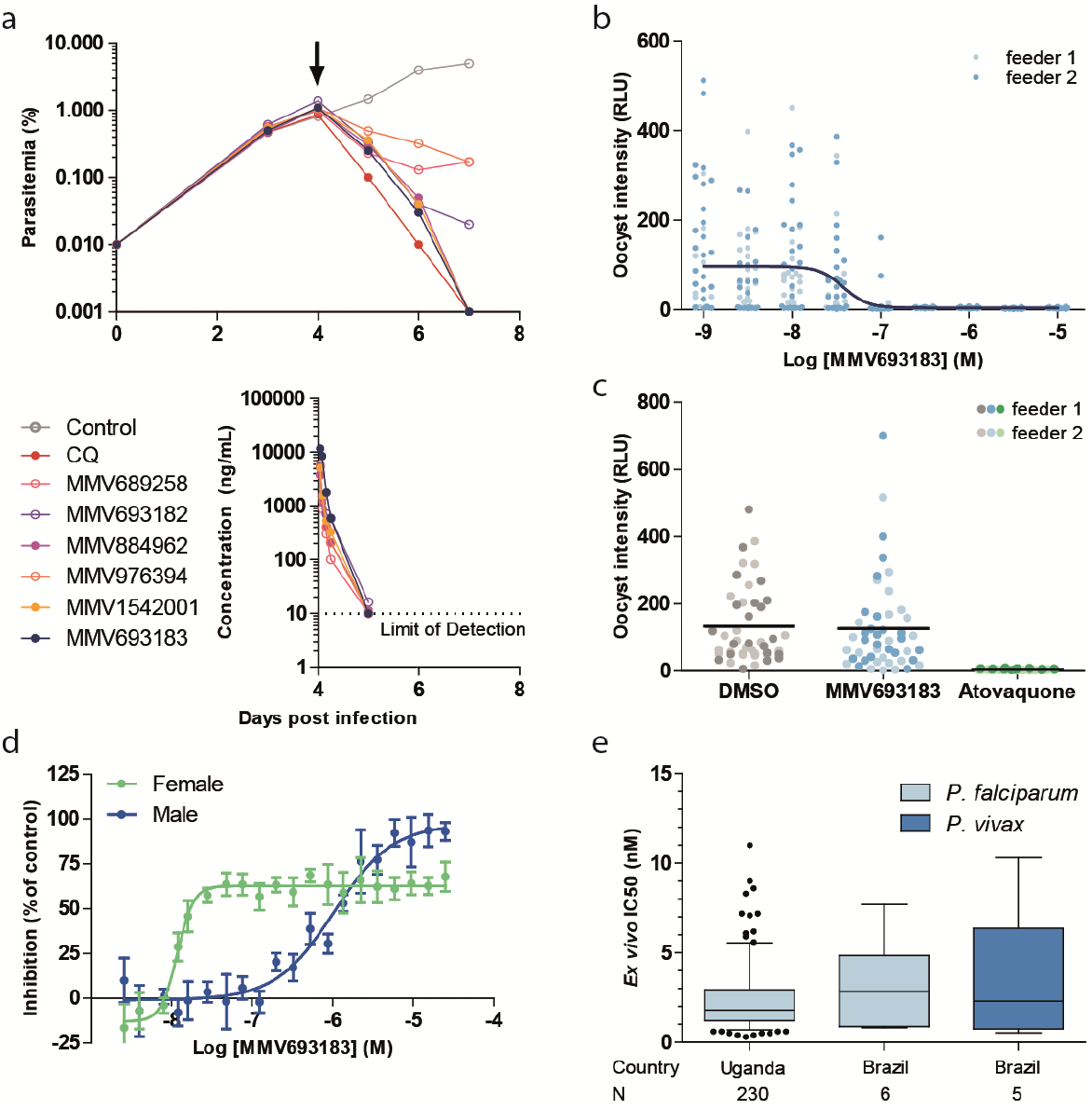
Antimalarial activity of the pantothenamide MMV693183. **a**, *In vivo* activity of novel pantothenamides. NSG mice were infected with *P. falciparum* on day 0. Mice were treated with pantothenamides by oral gavage (50mg/kg) (N = 2/compound) on day 4 (arrow) and parasitemia was quantified every day from day 3 onwards (top panel). The corrected concentration of pantothenamides in blood is indicated in the bottom panel. **b**, The activity of MMV693183 on *P. falciparum* (NF54-HGL) stage V gametocytes treated for 24 h before mosquito feeding in a single experiment with two replicates (feeder 1, 2). Oocyst intensity was measured by luminescence eight days after the feed. **c,** C) Oocyst intensities in mosquito midguts when *P. falciparum* (NF54-HGL) stage V gametocytes were exposed to 1 μM MMV693183, 100 nM atovaquone, or 0.1% DMSO within the mosquito blood meal. Oocyst intensity was quantified by luminescence eight days after feeding in a single experiment with two replicates (feeder 1, 2). **d,** Dual gamete formation assay upon treatment of female or male gametocytes with MMV693183 in four independent experiments (±SD). Typically, 150-250 exflagellation centers or 2000-3000 female gametes per field were recorded in the negative controls. **e,** *Ex vivo* activity of MMV693183 against field isolates of *P. falciparum* from Uganda (N = 230) in a parasite growth assay and against field isolates of *P. falciparum* (N = 6) and *P. vivax* (N = 5) from Brazil in a schizont maturation assay. Median IC_50_ values and 5-95 percentile are shown in a Box-Whisker plot. CQ, chloroquine.

**Table 1.**
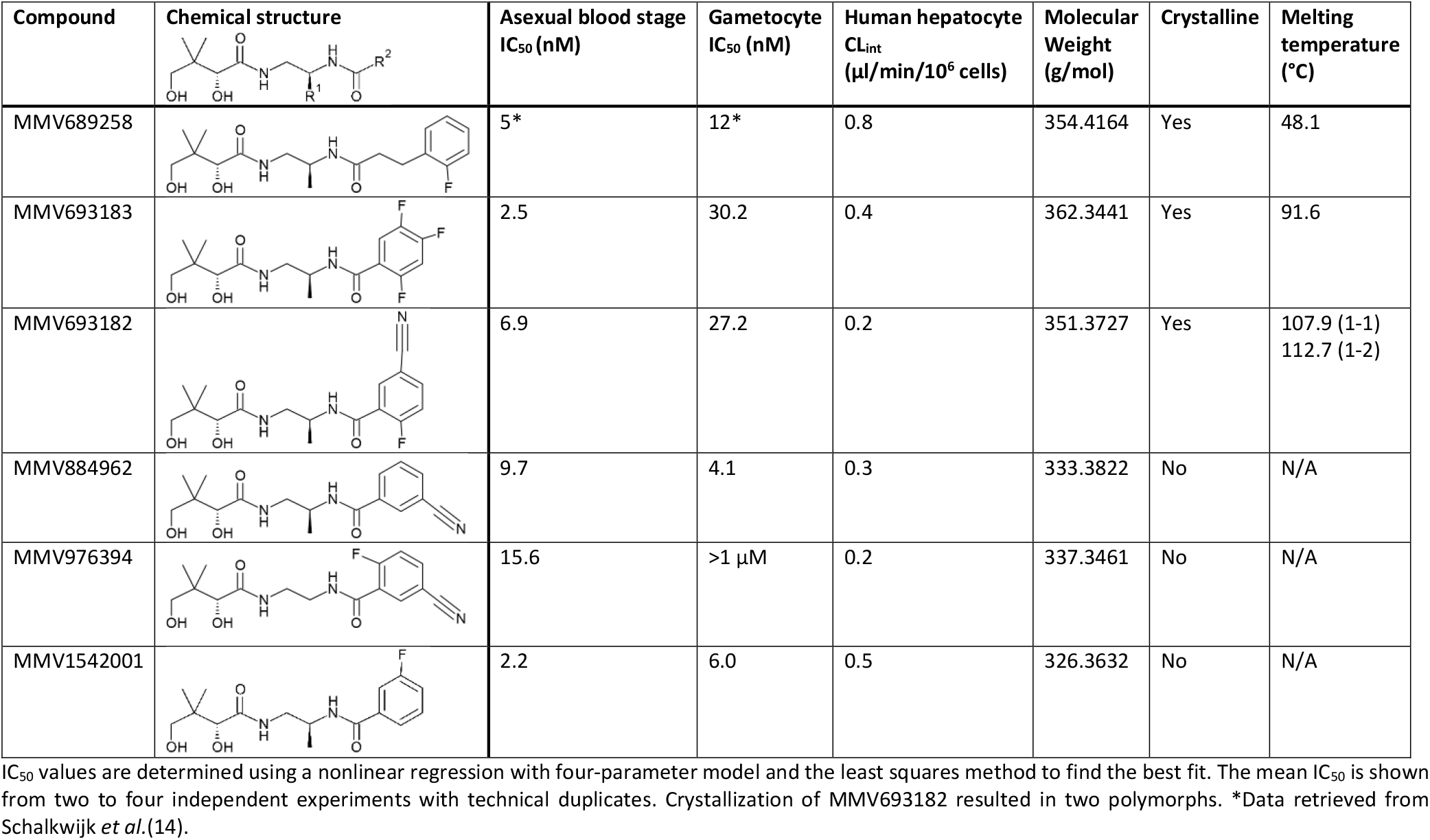
Physicochemical characteristics and *in vitro* activities of pantothenamides.

| Compound | Chemical structure | Asexual blood stage<br>IC<sub>50</sub> (nM) | Gametocyte<br>IC<sub>50</sub> (nM) | Human hepatocyte<br>CL<sub>int</sub><br>(μl/min/10<sup>6</sup> cells) | Molecular<br>Weight<br>(g/mol) | Crystalline | Melting<br>temperature<br>(°C) |
| --- | --- | --- | --- | --- | --- | --- | --- |
| MMV689258 |  | 5* | 12* | 0.8 | 354.4164 | Yes | 48.1 |
| MMV693183 |  | 2.5 | 30.2 | 0.4 | 362.3441 | Yes | 91.6 |
| MMV693182 |  | 6.9 | 27.2 | 0.2 | 351.3727 | Yes | 107.9 (1-1)<br>112.7 (1-2) |
| MMV884962 |  | 9.7 | 4.1 | 0.3 | 333.3822 | No | N/A |
| MMV976394 |  | 15.6 | >1 μM | 0.2 | 337.3461 | No | N/A |
| MMV1542001 |  | 2.2 | 6.0 | 0.5 | 326.3632 | No | N/A |

IC<sub>50</sub> values are determined using a nonlinear regression with four-parameter model and the least squares method to find the best fit. The mean IC<sub>50</sub> is shown from two to four independent experiments with technical duplicates. Crystallization of MMV693182 resulted in two polymorphs. \*Data retrieved from Schalkwijk *et al.*(14).

Asexual blood-stage parasites treated with MMV693183 in a parasite reduction rate (PRR) assay showed rapid killing activity. Parasitemia was reduced within 24 h and to below the detection limit within 48 h (Fig S3). This profile is similar to artemisinins (22) that constitute the fastest-acting class of clinical antimalarials available to date. MMV693183 was not efficacious against liver stages (Fig S4), similar to previous findings (14). Treatment of gametocytes 24 h before feeding to *A. stephensi*, inhibited oocyst formation with an IC_50_ value of 38 nM (Fig 1b), but treatment with 1 μM directly at the time of the mosquito feeds did not inhibit midgut infection (Fig 1c), confirming the gametocytocidal mode of action. MMV693183 specifically inhibited female gametocyte activation with an IC_50_ value of 12 nM, whereas male gamete formation was inhibited much less with an IC_50_ value of 1 μM (Fig 1d). We also performed *ex vivo* activity assays against *P. falciparum* and *P. vivax* field isolates from Uganda and Brazil. Encouragingly, all isolates were sensitive to the drug, with low nanomolar IC_50_ values (Fig 1e).

### A role for ACS in the mode of action of MMV693183

*In vitro* evolution and whole-genome analysis (IVIEWGA) experiments identified a single point mutation in *ACS* (PF3D7_0627800) resulting in a T648M amino acid change (dT648M) in *P. falciparum* Dd2-B2 and NF54 strains pressured at sublethal concentrations of MMV693183 (Table S3 and S4). Exposing different inocula of Dd2-B2 parasites to the compound identified a minimum inoculum of resistance of 1×10^9^ Dd2 parasites (Table S3, Fig S5). The resistant parasites showed a 13-77× IC_50_ shift against MMV693183 in an asexual blood-stage growth assay and also resulted in resistant gametocytes (>50× IC_50_ shift) (Fig 2a) that were able to transmit to mosquitoes (Fig S6a-c). CRISPR-Cas9 engineering of this mutation in wild-type parasites (cT648M) confirmed the resistance phenotype observed for the T648M mutation (cT648M) in both asexual and sexual blood stages (49× and >50× IC_50_ shift, respectively), and conferred cross-resistance to other PanAms (Fig 2a, Fig S7a-b). In addition, mutant parasite with a previously described T627A mutation in ACS (14) were also resistant to MMV693183 (Fig 2a). Metabolomic profiling showed that MMV693183 was converted into three CoA-precursors (Fig 2b, bottom panel) and reduced acetyl-CoA and 4-phosphopantothenate levels in infected RBCs in a dose-dependent manner (Fig 2b, upper panel). This is in line with previous observations of PanAm antimetabolite generation and suggests inhibition of ACS function.

**Fig 2.**
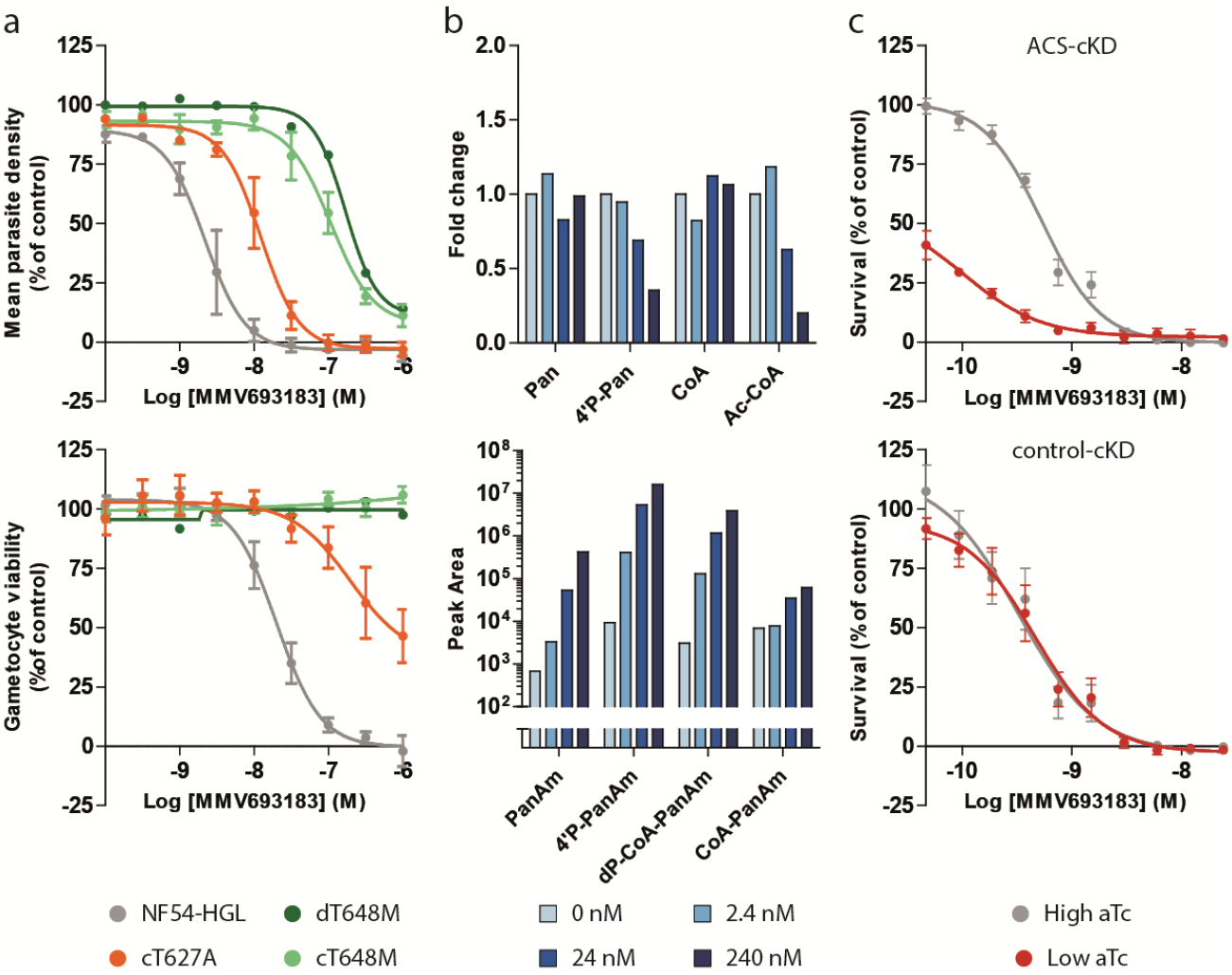
Role of ACS in the mode of action of MMV693183. **a,** Drug-sensitivity profiles with asexual (upper panel) or sexual (lower panel) blood-stage parasites without a mutation (NF54-HGL) or parasites with a T648M or T627A mutation in ACS. A MMV693183-induced resistant parasite line (dT648M) was tested in one experiment with two technical replicates and the CRISPR-engineered parasites (cT648M and cT627A) were tested in three independent experiments (two technical replicates per experiment). The average value for mean parasite density relative to controls ± SEM are shown. **b,** Concentration-dependent changes in levels of endogenous metabolites (upper panel) and pantothenamide antimetabolites (lower panel) upon treating *P. falciparum*-infected RBCs with MMV693183 or no drug. 3D7 parasites were synchronized at the trophozoite stage and treated with increasing concentrations of compound for 2.5 h and (anti)metabolites were quantified in two independent experiments with three technical replicates. Untreated parasites represent the background levels of MMV693183 metabolites. CoA could not be identified in the second experiment, therefore, only data from the first experiment are shown for the CoA level. The fold change is determined relative to no drug control (0 nM). Pan: pantothenate; 4’P-Pan: 4’-phosphopantothenate; Ac-CoA: acetyl-CoA. **c,** Drug-sensitivity assays on conditional knockdown parasites of ACS (upper panel) or a control target (lower panel) on asexual blood stages at low or high aTc (N = 3). The graphs show parasite survival based on a luminescence readout compared to controls ± SEM. aTc, anhydrotetracycline.

To further confirm a role for ACS in the mechanism of action to MMV693183, we used an ACS or control conditional knockdown parasite line (R. Summers, C.F.A. Pasaje, J.C. Niles, A.K. Lukens, submitted) using the TetR-DOZI system that can repress translation of the target gene when the drug anhydrotetracycline (aTc) is removed (23, 24). We cultured conditional knockdown ACS parasites (ACS-cKD) and control knockdown parasites (control-cKD) in low aTc (1.5 or 0 nM aTc, respectively) or high aTc (500 nM) conditions and exposed them to different doses of MMV693183. The IC_50_ for the ACS-cKD parasite line decreased 5-fold upon knockdown conditions (low aTc), showing hypersensitivity of these parasites to the compound, while there was no difference in sensitivity for the negative knockdown control (Fig 2c). This supports our hypothesis that PanAms act in an ACS-dependent manner.

### CoA-PanAm targets ACS

To provide conclusive evidence that PanAms directly bind to ACS, thereby inhibiting ACS activity, we used the recently developed cellular thermal shift assay (CETSA) (25, 26) to test the thermal stability of ACS upon treatment with PanAms. Rabbits were immunized with a recombinant ACS fragment and the induced anti-serum was used to detect ACS (Fig S8). Following incubation of asexual blood stages with compounds for 1 h, neither the parent compound MMV693183 nor its derivative 4’P-MMV693183 affected ACS stability. However, the CoA-MMV693183 metabolite clearly stabilized ACS upon temperature increase (Fig 3a). This supports the notion that PanAms form active CoA-PanAm antimetabolites that target ACS. To provide further evidence that CoA-PanAm targets ACS, we immunopurified ACS from parasite lysates and established an ACS activity assay. CoA-MMV693183 inhibited ACS activity with an IC_50_ of 154 nM, whereas 4’P-MMV693183 only showed weak activity (Fig 3b). This shows for the first time that the CoA-PanAm is the active metabolite that inhibits ACS.

**Fig 3.**
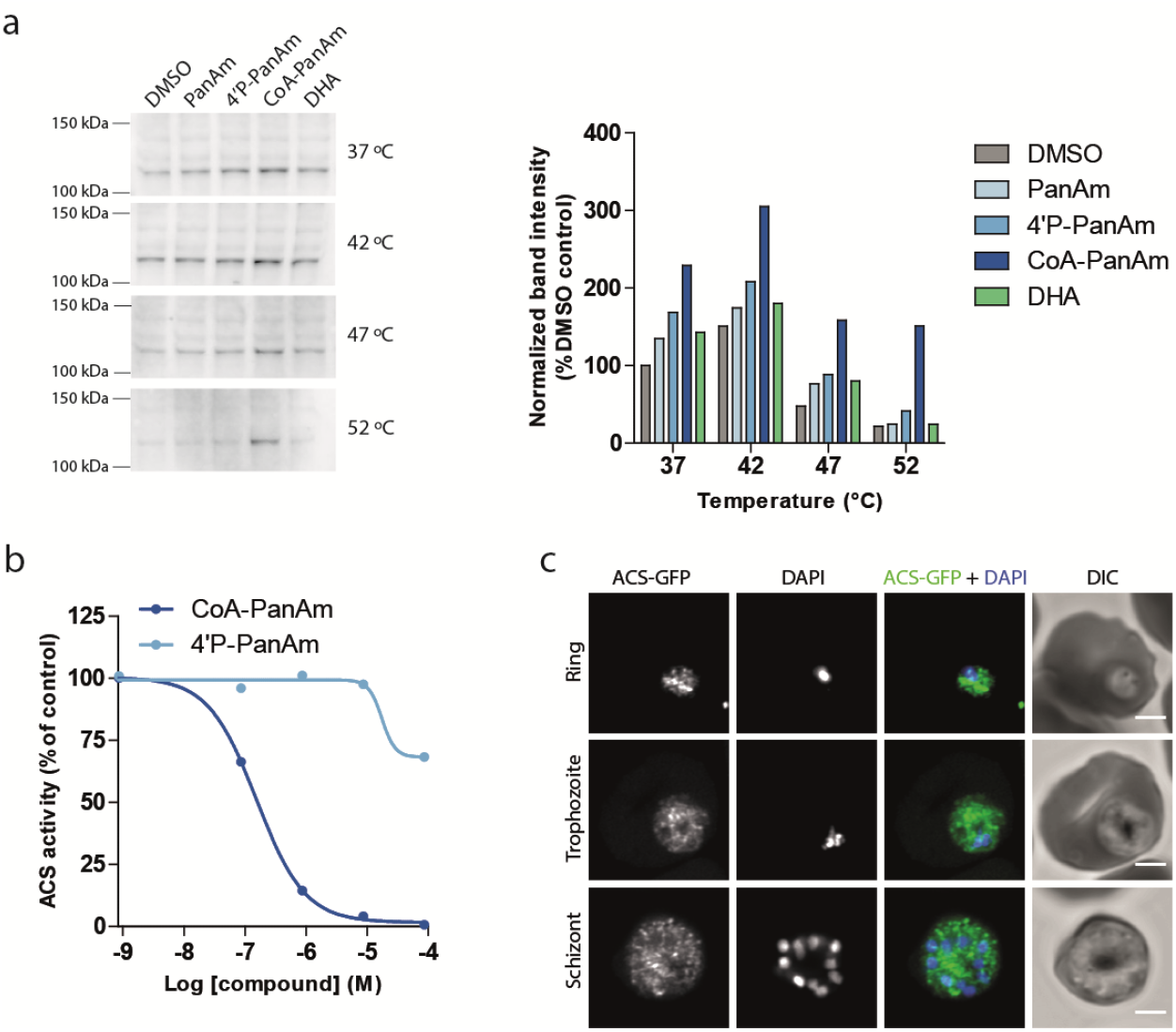
CoA-PanAm binds to and inhibits ACS. **a,** Cellular thermal shift assay on *P. falciparum* lysate. Parasite lysate at 2.1 mg/ml was aliquoted and treated with 1 μM MMV693183 or metabolites thereof, or with DHA (negative control) for 30 min, followed by a 3-min incubation at different temperatures (N = 1). Protein stabilization was analyzed on a western blot (left panel) and band intensities (ACS molecular weight = 113.8 kDa) were quantified and normalized to DMSO treatment at 37°C (right panel). **b,** ACS activity in a dose-response assay. ACS was immunopurified from parasite lysate using rabbit immune serum. The activity was measured using ^14^C-labeled sodium acetate upon treatment with metabolites of MMV693183 and normalized to the no drug control (N = 2). **c,** Immunofluorescence microscopy of parasites with ACS fused to GFP. Depicted are representative images of asexual blood-stage parasites stained with anti-GFP antibodies and DNA stained with DAPI. Scale bars, 2 μm. PanAm: MMV693183; 4’P-PanAm: 4’P-MMV693183; CoA-PanAm: CoA-MMV693183.

ACS is predicted to provide acetyl-CoA for a variety of processes in the parasite, including fatty acid elongation in the endoplasmic reticulum and post-translational modifications in the cytosol and nucleus (27–29). To begin to explore whether inhibition of ACS could affect these downstream pathways, we studied the localization of ACS using an endogenous GFP-tagged ACS parasite line (Fig S9a-b). ACS-GFP localized to the cytosol, although it was unclear whether it is also expressed in the nucleus in *P. falciparum* (Fig 3c), as previously observed in apicomplexan parasites (27, 30, 31). To verify this localization, we stained wild-type parasites with ACS immune serum. A possible perinuclear or cytoplasmic signal was observed, but we could not detect an evident nuclear signal (Fig S10).

### MMV693183 safety

Given that CoA metabolism clearly plays a central and crucial role in human cells, it was important to examine whether MMV693183, like MMV689258 (14), acts selectively and specifically on the parasite without affecting the human host. Our studies revealed that treatment with MMV693183 was not cytotoxic to either HepG2 cells or primary human or rat hepatocytes, did not affect human cardiac ion channels, including the Kv11.1 (hERG) channel, did not induce or modulate CYP activity and was negative in AMES and micronucleus tests (Table S5). In addition, MMV693183 did not show cross-reactivity to a panel of human receptors, enzymes, or channels. In all assays, inhibition was <50% at a test concentration of 10 μM (Table S5). A UV-scan did not reveal a liability for phototoxicity as there was no detectable absorption above 290 nm (Table S5). Unlike the prophylactic antimalarial drug primaquine, MMV693183 did not show signs of hemolytic toxicity in a mouse model of human G6PD deficiency (Fig S11) (32).

Preliminary *in vivo* safety of MMV693183 was tested in rats with a seven-day repeat dose study (Table S6) focusing on glucose, triglycerides and urea, as these or intermediates thereof were previously shown to be affected upon disruption of the CoA pathway in mice (33). The mean C_max_ (range of 35.5 to 501 μM) and AUC (41.2 to 1080 h*μM) were mostly dose-proportional in males and females on day 1. However, the exposure (mean range C_max_: 15.6 to 168 μM; mean range AUC: 11.4 to 168 h*μM) on day 7 was lower than on day 1, with a decrease of up to 87% in AUC, except for females at 60 mg/kg/day, in which the exposure was similar (Table S7). Body weight was unchanged (Table S8), and no mortality or clinical signs of toxicity occurred during the observation period. No significant differences were observed for glucose, triglycerides or urea concentrations (Fig 4a-c). Other clinical chemical parameters were mostly unaffected or showed only minor effects (Table S9). This highlights that MMV693183 does not induce the drastic effects demonstrated previously in rodents upon chemical disruption of the CoA pathway by hopantenate (33).

**Fig 4.**
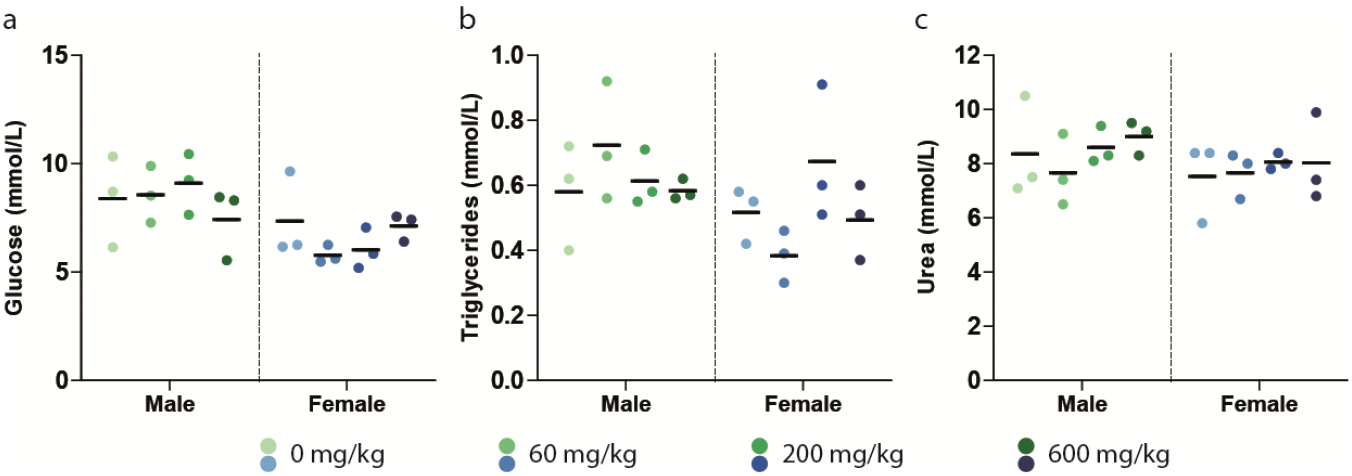
*In vivo* safety of MMV693183. **a-c,** Evidence of i*n vivo* toxicity was examined in male and female Wister Han rats (N = 3 per condition) treated for seven days with MMV693183. Glucose (**a**), triglycerides (**b**) and urea (**c**) concentrations were measured in rats (male or female) treated with different doses of MMV693183. Significance was determined using One-Way Anova with the Bonferroni’s Multiple Comparison Test.

### Pharmacokinetic properties

Pharmacokinetic (PK) studies were performed in order to support a human dose prediction. The MMV693183 PK profiles in mice, rats and dogs were examined using two-compartment models that were fit to plasma concentration-time data observed after oral and intravenous dosing (Fig S12, Table S10-12). Even though the Caco-permeability assay suggested moderate absorption and active efflux *in vitro* (Table S13), MMV693183 was absorbed rapidly (Tmax of 0.5 h) *in vivo* and had an excellent oral bioavailability (64% in dogs to 121% in rats) (Table S10-12). The total clearance was 13.4, 21.1 and 11.5 ml/min/kg, and the half-life was 1.2, 3.1 and 4.2 h in mice, rats and dogs, respectively (Table S10-S12). The plasma protein binding was overall low (ranging from 33% in mouse to 52% in human plasma) (Table S14) and blood to plasma ratio ranged from 0.91 to 0.98. The main route of metabolism in an *in vitro* human hepatocyte relay assay was an oxidation, followed by a glucuronide conjugation and dehydrogenation of the hydroxyl groups (Table S15). In plasma and urine collected from dogs, we also detected oxidized, dehydrogenized and glucuronide-conjugated metabolites (Table S16). In rats, 36-40% of the drug was eliminated in urine, while in dogs this was predicted to be 8.9%, excluding values from two dogs with only <20 ml urine (Table S17-18). Metabolic stability of MMV693183 was assessed in primary hepatocytes from mice, rats and dogs. The observed values correlated well with the non-renal clearance observed *in vivo*, suggesting that the non-renal clearance is mainly via a hepatic route (Table S19). For mice, no renal clearance data were available, but the total observed *in vivo* clearance amounted to 13.4 ml/min/kg whereas the predicted hepatic clearance was 7 ml/min/kg. This implies that 6.4 ml/min/kg (48%) of total clearance was contributed by the kidney, in line with the proportion of renal clearance in rats.

### Human pharmacokinetic predictions

Two approaches were considered to predict human clearance. First, a clearance exponent of 0.967 was derived using simple allometry, which was higher than typical ranges for this parameter (0.67-0.75) (34). Consequently, the maximum life-span potential (MLP) correction was implemented resulting in an estimated total clearance of 1.8 ml/min/kg (35). Second, human hepatic clearance was measured at 0.51 ml/min/kg based on the *in vitro* hepatocyte clearance of 0.4 μl/min/10^6^ cells. Human renal clearance was predicted at 0.60 ml/min/kg based on renal clearance in dogs (Table S20) corrected for plasma protein binding (52%; Table S14) and kidney blood flow, which was previously shown to be a good predictor (36). Given the excellent correlation between the *in vitro* and *in vivo* data for the animal studies (Table S19), the *in vitro* prediction of 0.51 ml/min/kg was preferred to predict total human clearance. This yielded a total clearance of 1.11 ml/min/kg, and a predicted human half-life of 32.4 h. Further human PK parameters were predicted using allometric scaling (Fig S13). Based on the Caco-2 permeability and thermodynamic solubility data (Table S2, S13), the bioavailability was predicted to be 96% (GastroPlus). The predicted human parameters are shown in Table S20.

### Prediction of the human efficacious dose using a PKPD model

*In vivo* efficacy data from three female NSG mice studies were pooled to evaluate the PKPD relationship of MMV693183 and derive key PD parameters such as MIC and MPC_90_ (37). For all single dose groups, the concentration of MMV693183 decreased to below the *in vitro*-determined IC_99_ (36.1 nM) corrected for its free fraction (67% in mice) within 24 h (Fig S14), while parasites were being cleared at four to six days. However, parasites recrudesced within 15 days after treatment (Fig S15). The PK and PD profiles were well captured by a three-compartment PK model with zero-order absorption and a linear elimination, and the *in vitro* clearance PD model, respectively (Table S21 and S22; Fig S14, S15). Differences in the initial decline of parasitemia between NSG mice studies were captured in the model explaining part of the parasitemia clearance parameter interindividual variability. The PKPD model predicted a MIC and MPC_90_ of 3.2 and 38.4 ng/ml, respectively.

The human dose prediction to achieve a 9 log total parasite reduction was predicted at 10 and 20 mg using the total clearance values from *in vitro* and allometric prediction, respectively (Fig 5a). To achieve a 12 log total parasite reduction, the predicted dose was 15 and 30 mg, respectively (Fig 5a, Table S23). A local sensitivity analysis was performed on total clearance and EC_50_ to evaluate the impact of the variation of both parameters on the human efficacious dose prediction. The most sensitive parameter for dose prediction was total clearance. With a total clearance of 5-fold higher than the predicted value using the *in vitro* hepatocyte clearance approach, the dose predicted to achieve 12 log total parasite reduction would be close to 625 mg for an adult (9 mg/kg) (Fig 5b), suggesting that MMV693183 could still be a valid candidate for a single dose cure.

**Fig 5.**
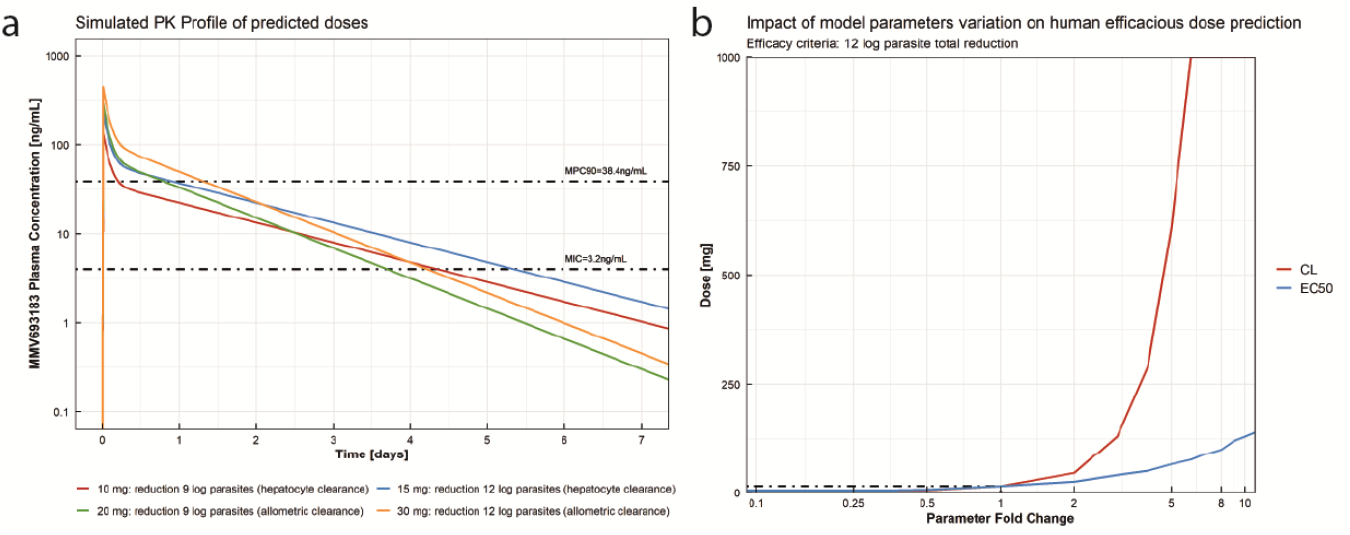
Human efficacious dose prediction. **a,** MMV693183 plasma concentration after the predicted efficacious human doses of 10, 15, 20 and 30 mg according to the efficacy criteria and human clearance prediction method. **b,** Local sensitivity analysis of the impact of total clearance and EC_50_ variation on the estimated efficacious dose, defined by a 12 log total parasite reduction efficacy criteria. LLoQ: Lower Limit of Quantification.

## Discussion

Following an extensive chemical optimization process, we have identified the novel compound PanAm MMV693183, which has low nanomolar potency against asexual blood stages of both *P. falciparum* and *P. vivax*, and against *P. falciparum* gametocytes. We showed its favorable physicochemical properties with potential to be developed into a single dose malaria cure. Furthermore, we revealed that the antimetabolite, CoA-MMV693183 acts upon ACS, thereby targeting an unexplored pathway for antimalarial therapy. These promising characteristics of MMV693183 support the recent selection of this drug for continued (pre)clinical development (4).

While pantothenate analogs have long been explored, stable and highly potent pantothenamides against *P. falciparum* have only been developed in the last decade (8, 11–13), but MMV693183 is the first pantothenamide to meet the criteria for further (pre)clinical development. This compound has improved *in vitro* and *in vivo* potency, metabolic stability, and a prolonged predicted human half-life compared to previously synthesized pantothenamides (12–14). Its promising potency is reflected in the predicted human efficacious single dose of 100 mg for treatment of clinical malaria. Furthermore, MMV693183 is highly potent against *P. vivax*, another major contributor to the malaria burden (1), supporting the activity of PanAms against multiple species also including *P. knowlesi* (38). The importance of combining MMV693183 with a partner drug is highlighted by the possibility of generating resistance against this compound that can also be transmitted to the mosquito vector. Further research to identify the ideal partner drug is needed. However, we could envision a drug combination that is able to target pantothenamide-resistant gametocytes.

The identified mutation in *ACS*, but not in *ACS11*, in MMV693183-resistant parasites, supported our previous hypothesis that ACS is primarily targeted by metabolized PanAms, although definitive proof was lacking (14, 17, 18). Here, we have conclusively demonstrated for the first time that CoA-PanAm is the active metabolite that inhibits ACS. With the previously suggested role of ACS in regulating the acetylome, transcriptome and metabolome (including fatty acid elongation) (27, 28, 30, 31, 39) and the corresponding cytoplasmic/perinuclear (this study) and nuclear localization of ACS for these functions (31), it could be hypothesized that these pathways are affected by CoA-PanAms. Similar to PanAms, inhibitors of histone deacetylases or acetylases (HDAC or HAT), enzymes that regulate histone acetylation, have dual-stage activity targeting both asexual and sexual blood-stage parasites (40–42). The marked difference between female and male sexual stage activity of MMV693183 could be related to one of the possible downstream consequences of ACS inhibition that may be more important in female than in male gametocytes. Alternatively, the differential activity against male gametocytes compared to females may be explained by lower availability of PanAm; for example through reduced uptake, increased export or increased breakdown. While the target of PanAms has been identified, the further downstream consequences are not yet understood.

It is clear that PanAms target a central pathway of *Plasmodium* parasites, which is conserved among many eukaryotes and prokaryotes. Previous studies on hopantenate, a compound that affects CoA metabolism, shows lethal toxicity within 15 days, a significant reduction in glucose and altered liver metabolism in mice (33). It is therefore of utmost importance to test the safety of PanAms. In a preliminary safety study, MMV693183 did not reduce glucose, and was not lethal to rats within the 7 day-period. This was in contrast with hopantenate treatment, although these mice were on a pantothenate-free diet (33). Furthermore, MMV689258 does not affect CoA metabolism in primary hepatocytes (14). Comprehensive toxicology studies are the subject of ongoing research and are critical for determining the safety of this compound. Even though no off-target activity was identified in our study, a cautionary note could be the weak effect on HDAC11, which is in line with the mechanism of action of MMV693183. This will be scrutinized further in ongoing toxicology studies.

A few limitations of our PKPD model need to be considered, which could affect the final dose predictions. The PK model based on humanized mouse data showed a high relative standard error for a few of the estimated parameters, which may lead to uncertainties in the PD parameters used for the final dose predictions, such as the EC_50_. However, the sensitivity analysis showed a limited impact of the variation of EC_50_ on the dose predictions, identifying MMV693183clearance in humans as the most sensitive parameter. Despite this, dose predictions using human clearance predicted from allometry (worst-case scenario) and sensitivity analysis of potential human clearance variations yielded encouraging results for MMV693183.

In conclusion, we provide a new preclinical candidate MMV693183 that is a promising multi-stage active compound and that acts on a pathway that is not currently targeted by clinical antimalarials. This agent has the potential to be developed into a single dose cure, and upon successful development may therefore aid in ongoing efforts to achieve malaria elimination.

## Materials

### Ethics statement

Animal experiments performed at The Art of Discovery (TAD) were approved by The Art of Discovery Institutional Animal Care and Use Committee (TAD-IACUC). This committee is certified by the Biscay County Government (Bizkaiko Foru Aldundia, Basque Country, Spain) to evaluate animal research projects from Spanish institutions according to point 43.3 from Royal Decree 53/2013, from the 1^st^ of February (BOE-A-2013-1337). All experiments were carried out in accordance with European Directive 2010/63/E.

The animal experiments carried out at the Swiss Tropical and Public Health Institute (Basel, Switzerland) were adhering to local and national regulations of laboratory animal welfare in Switzerland (awarded permission no. 2303). Protocols are regularly reviewed and revised following approval by the local authority (Veterinäramt Basel Stadt).

Aptuit is committed to the highest standards of animal welfare and is subject to legislation under the Italian Legislative Decree No. 26/2014 and European Directive No. 2010/63/UE. Animal facilities are authorized by the Italian Ministry of Health with authorization n. 23/2017-UT issued on 29th November 2017 according to art. 20 of Legislative Decree No. 26/2014. Furthermore, general procedures for animal care and housing are in accordance with the Association for Assessment and Accreditation of Laboratory Animal Care (AAALAC) recommendations.

Animal procedures to determine the hemolytic toxicity were approved by the University of Colorado Anschutz Medical Campus Institutional Animal Care and Use Committee.

All animal studies had the approval of the Institutional Animal Ethics Committee (IAEC) of TCG Lifesciences Pvt. Ltd and were conducted in accordance with the guidelines of the Committee for the Purpose of Control and Supervision of Experiments on Animals (CPCSEA), Government of India.

The seven-day repeat dose study in rats was reviewed and agreed by the Animal Welfare Body of Charles River Laboratories Den Bosch B.V. within the project license AVD2360020172866 approved by the Central Authority for Scientific Procedures on Animals (CCD) as required by the Dutch Act on Animal Experimentation (December 2014).

More information on animal experiments can be found in Table S24.

For collection of blood for *ex vivo* activity studies in Brazil and Uganda, all participants or their parents/guardians signed a written informed consent before blood collection. Patients were promptly treated for malaria after blood collection, following national guidelines. The *ex vivo* activity study in Brazil was approved by the Ethics Committee from the Centro de Pesquisa em Medicina Tropical - CEPEM-Rondônia (CAAE 61442416.7.0000.0011). The *ex vivo* activity study in Tororo, Uganda was approved by the Makerere University Research and Ethics Committee, the Uganda National Council for Science and Technology, and the University of California, San Francisco Committee on Human Research.

The human biological samples were sourced ethically and their research use was in accord with the terms of the informed consents under an IRB/EC approved protocol.

### Chemistry

The synthesis of the PanAms is included in patent application EP3674288A1 (20), which states: “Characteristic features of the analogs concern the moieties flanking the inverted amide; the carbon atom flanking the inverted amide in the center portion of the molecule could comprise a methyl substituent, the two nitrogen atoms are separated by a linker of two carbon atoms, and the moiety flanking the inverted amide at the distal portion of the molecule is a (hetero)aromatic, optionally substituted, ring or ring system, bonded directly to the carbonyl group of the inverted amide.”

Crystal screening for each further characterized pantothenamide was performed by a commercial service (Crystal Pharmatech Co., Ltd.) under 36 conditions using a variety of crystallization methods, including liquid vapor diffusion, slow evaporation, slurry conversion and salt/co-crystal formation and solvents. X-ray power diffraction (XRPD) patterns were collected by Bruker X-ray powder diffractometers.

### Parasite culture and *in vitro* efficacy of pantothenamides

The *P. falciparum* strains Dd2-B2 (a clone of Dd2), 3D7, NF54 and the luminescent-reporter strain NF54-HGL (43) were cultured in RPMI 1640 medium supplemented with 25mM HEPES, 382 μM hypoxanthine, 26 mM NaHC0_3_, 10% human blood type A serum or 0.5% AlbuMAX II, and 3-5% human blood type O red blood cells (RBCs) (Sanquin, the Netherlands) at 37°C in 3% O_2_, 4% CO_2_.

Replication assays were performed using a SYBR Green method as described previously (44). Briefly, parasites were diluted to 0.83% parasitemia, 3% hematocrit in 30 μl medium and added to 30 μl of diluted compounds in medium (0.1% DMSO final concentration) in black 384-wells plates. After a 72-h incubation, 30 μl of SYBR Green diluted in lysis buffer was added according to the manufacturer’s protocol (Life Technologies). Fluorescence intensity was measured on a BioTek Synergy 2 Plate Reader after 1-h incubation and was normalized to a DMSO control (100% growth) and DHA- or epoxomicin-treatment (no growth). To define the IC_50_ of MMV693183-resistant Dd2-B2 parasites, ring-stage cultures at 0.3% parasitemia and 1% hematocrit were exposed for 72 h to a range of concentrations of MMV693183 along with drug-free controls. Parasite survival was assessed by flow cytometry on an Accuri C6 (BD Biosciences) using SYBR Green and MitoTracker Deep Red FM (Life Technologies) as nuclear stain and vital dye, respectively. To assess the effect of conditionally perturbing ACS expression and treatment with MMV693183 on parasite growth, synchronous ring-stage ACS conditional knockdown parasites (ACS-cKD) or a control conditional knockdown line (control-cKD; previously generated parasites with a yellow fluorescent protein with regulatory TetR aptamers in the 3’untranslated region (UTR) integrated in the *cg6* chromosomal locus (23)) were cultured in high (500 nM) and low (1.5 or 0 nM, respectively) concentrations of anhydrotetracycline (aTc) and incubated with serially diluted MMV693183. Luminescence was measured after 72 h using the Renilla-Glo(R) Luciferase Assay System (Promega E2750) and the GlomAX® Discover Multimode Microplate Reader (Promega). The luminescence values were normalized to DMSO vehicle (100% growth) and 200 nM chloroquine-treated (no growth) samples as controls.

The antimalarial killing rate was determined by GlaxoSmithKline (GSK, Tres Cantos, Madrid, Spain) as described previously (22). Briefly, 0.5% 3D7 (BEI Resources) *P. falciparum* parasites (≥80% ring-stage population) at 2% hematocrit were treated with 10x IC_50_ of MMV693183 (40 nM in 3D7 parasites) or pyrimethamine (0.94 μM) for 120 h and drug was renewed daily. Parasite samples were taken every 24 h and drug was washed out, followed by four independent, 3-fold serial dilutions in 96-wells plates. The number of viable parasites was determined on day 21 and 28 by counting the wells with parasite growth. Parasite growth was measured by uptake of ^3^H-hypoxanthine in a 72-h assay and was back-calculated to viable parasites using the following equation X^n-1^ where n is the number of parasite-positive wells and X the dilution factor.

Gametocyte viability assays on NF54-HGL parasites were performed using an adapted high-throughput protocol as previously described (14, 45). In short, asexual blood-stage parasite cultures were set up at 1% parasitemia in a semi-automated shaker system at 5% hematocrit (46). From day four until day eight or nine, parasites were treated with 50 mM *N*-acetyl glucosamine to eliminate all asexual blood-stage parasites. Subsequently, gametocytes were isolated by a Percoll density gradient centrifugation (45). At day 11, gametocytes were seeded (5,000 per well) in 30 μl in white 384-well plates containing 30 μl of compounds diluted in medium (0.1% DMSO). After a 72-h incubation, 30 μl of ONE-Glo reagent (Promega) was added according to manufacturer’s protocol and luminescence was quantified using the BioTek Synergy 2 Plate reader. Values were normalized to DMSO- and epoxomicin-treated controls.

Activity of MMV693183 against female and male gametocytes was assessed in a dual gamete formation assay (DGFA) as described previously (47). Briefly, mature *P. falciparum* NF54 gametocyte cultures were added to 384-well plates containing DMSO or different concentrations of MMV693183 (in <0.25% DMSO) or Gentian Violet (12.5 μM). After a 48-h incubation, gamete formation was stimulated by a drop in temperature (from 37 °C to 26 °C), and the addition of xanthurenic acid (2.5 μM). At 20 min after induction, exflagellation was recorded by automated time-lapse microscopy. After data collection, the plate was returned to a 26°C incubator and incubated for 24 h. Female gamete formation was assessed by live staining with a Cy3-labelled anti-Pfs25 monoclonal antibody and recorded by automated microscopy.

To assess parasite development in hepatocytes, cryopreserved human primary hepatocytes (Tebu-Bio lot: HC10-10) were thawed according to the manufacturer’s protocol and seeded (50,000 cells per well) in collagen-coated 96-well plates (Greiner). Cells were cultured at 37°C in 5% CO_2_ and medium was refreshed after 3 h and 24 h. Salivary glands from *Anopheles stephensi* mosquitoes were dissected to obtain NF54 sporozoites that were added (60,000 per well) to hepatocytes 48 h post-thawing. Plates were spun down and sporozoites were incubated with hepatocytes for 3 h. Subsequently, sporozoites were aspirated and compounds diluted in hepatocyte medium, were added to the hepatocytes (0.1% DMSO final concentration). Medium containing compounds was refreshed daily for four days. Hepatocytes were fixed with ice-cold methanol and samples were blocked with 10% fetal bovine serum (FBS) in PBS. Samples were incubated with rabbit anti-HSP70 (1:75, StressMarq) in 10% FBS for 1-2 hours followed by incubation with secondary goat anti-rabbit AlexaFluor 594 antibody (1:1000, Invitrogen) in 10% FBS for 30 min. Samples were washed with PBS containing 0.05% Tween 20 between different steps. Cells were imaged on the Biotek Cytation and images were analyzed automatically using FIJI software.

### *In vivo* efficacy of pantothenamides

The effect of pantothenamides on *P. falciparum* Pf3D7^0087/N9^ (48) *in vivo* was assessed in female NSG mice (NODscidIL2Rγ^null^) at the Swiss Tropical and Public Health Institute (Basel, Switzerland) as described previously (Table S24) (14, 49). Briefly, humanized mice were engrafted daily with human erythrocyte suspensions from days −11 to day 6. After 11 days (day 0), mice were injected intravenously with 3×10^7^ infected RBCs in a volume of 0.1 ml. On day 4, groups of n = 2 mice were treated with a single dose pantothenamides or chloroquine (50 mg/kg) by oral gavage. The hematocrit of all dosed mice and an untreated control group (n = 4 mice) was determined by fluorescence-activated cell sorting and parasitemia was analyzed by microscopy on >10,000 RBCs as described before (50). Samples to quantify compound metabolites were collected and prepared at different time points (1, 2, 4, 6, and 24 h after treatment) for each mouse by mixing 20 μl of whole blood with 20 μl of Milli-Q, followed by immediate freezing of samples on dry ice. Samples were processed under protein precipitation methods and analyzed by LC-MS/MS for quantitation in a TSQ Quantum Access (Thermo Fisher Scientific, San Jose, CA, USA). The lower limit of quantification was 5 ng/ml.

Transmission-blocking activity of compounds was determined as described previously (51). Briefly, NF54-HGL parasites were set up at 1% parasitemia in a semi-automated shaker system at 5% hematocrit. After 14 days of culturing, stage V gametocytes were treated with a range of concentrations of compound for 24 h before feeding, or with 1 μM MMV693183 or 100 nM atovaquone directly upon feeding to *A. stephensi*. Eight days after the feed, luminescence was quantified to determine oocyst intensity.

### *Ex vivo* efficacy of pantothenamides

For *ex vivo* pantothenamide activity studies, patients with *P. falciparum* or *P. vivax* were recruited at the Research Center for Tropical Medicine of Rondonia (CEPEM) in Porto Velho (Brazilian Western Amazon). A schizont maturation assay was performed using parasites obtained from mono-infected patients. A total of 44 patients were recruited who did not use any antimalarial in the previous months and/or present with symptoms of malaria, but had a parasitemia between 2,000 and 80,000 parasites/μl. Isolates from patients were excluded if (i) <70% of parasites were rings at the time of sample collection (n = 11), (ii) no schizont maturation was observed (n=9), or if (iii) the number of inviable parasites in untreated control was higher than the number of maturated schizonts in the treated condition (n = 4), leaving 20 patients to be included. Peripheral venous blood (5 ml) was collected by venipuncture in heparin-containing tubes, plasma and the buffy coat were removed, RBCs were washed and subsequently filtered in a CF11 cellulose column. Blood was diluted to 2% hematocrit in either RPMI 1640 medium (*P. falciparum)* or McCoy’s 5A medium (*P. vivax)* supplemented with 20% compatible human serum. Parasites were incubated with MMV693183 at final concentrations ranging between 0.25 and 500 nM in a hypoxia incubator chamber (5% O_2_, 5% CO_2_, 90% N2). The incubation of parasites with compound was stopped when 40% of the ring-stage parasites reached the schizont stage (at least three distinct nuclei per parasite) in the untreated control wells. The number of schizonts per 200 asexual blood-stage parasites was determined and normalized to control. An assay was considered valid when compound was incubated with parasites for at least 40 h.

An *ex vivo* growth inhibition assay was performed on fresh clinical *P. falciparum* isolates in Uganda. Blood was collected from patients aged ≥6 months presenting to the Tororo District Hospital, Tororo District, or Masafu General Hospital, Busia District with clinical symptoms suggestive of malaria, Giemsa-stained thick smears positive for *P. falciparum* infection, and ≥0.3% parasitemia determined by Giemsa-stained thin smears. Up to 5 ml of blood was drawn by venipuncture from 109 participants at Tororo District Hospital and 121 at Masafu General Hospital. Parasites were diluted to 0.2% parasitemia in 2% hematocrit and incubated for 72 h with serial dilutions of MMV693183 (0.1% DMSO) in a 96-well microplate and stored in a humidified modular incubator (2% O_2_, 3% CO_2_, 95% N2). Parasite density was quantified by fluorescence after incubation with SYBR Green lysis buffer measured on a BMG Fluostar Optima plate reader, as previously described (52).

### *In vitro* safety and toxicity assays

Safety studies were performed by commercial services using their standard protocols. Off-target activities of 10 μM MMV693183 were investigated using binding, enzyme and uptake assays (Eurofins CEREP, Celle-Lévescault, France). Phototoxicity of MMV693183 was assessed by exposing the compound to different wavelengths. Genotoxicity was investigated using the Ames test (Bacterial Reverse Mutation Assay) (Covance Laboratories Ltd, North Yorkshire, England). *In vitro* mammalian cell micronucleus screening assay was studied in human peripheral blood lymphocytes (BioReliance Corporation, Rockville, USA). Cardiotoxicity was determined against the hERG channel in an automated patch clamp assay using the Qpatch or against the hNa_V_ 1.5, hK_V_ 1.5 and hCa_V_ 1.2 using a manual patch clamp technique (Metrion Biosciences, Cambridge, UK). CYP450 induction and cytotoxicity assays were performed by a commercial service (KaLy-Cell, Plobsheim, France) according to their standard protocols. The IC_50_ value of CYP1A2, CYP2D6, CYP3A4 and CYP2C19 inhibition was determined in duplicate by a commercial service (TCG Lifesciences) using their standard protocols.

### Exploratory *in vivo* safety and toxicology studies

Hemolytic toxicity was determined in female NSG mice (Jackson Laboratories) using erythrocytes from a G6PD-A-deficient blood donor (0.4 u/g hemoglobin). Mice were engrafted with 3.5×10^9^ human RBCs intraperitoneally for fourteen days to obtain >60% human RBCs. Mice were treated for four days with vehicle control (PBS) or MMV693183 (10, 25, 50 mg/kg), or primaquine (12.5 mg/kg) for three days. Spleen weight was quantified on day seven and hemolysis was assessed on day zero, four and seven, by quantifying human RBCs and murine reticulocytes on a Fortessa Flow Cytometer (BD BioSciences) using anti-glycophorin A-FITC and anti-CD71-FITC and anti-TER119-PE, respectively.

The preliminary maximum tolerated dose study (53) was undertaken at a commercial service (Charles River, ‘s-Hertogenbosch and Groningen, Netherlands). Briefly, Wistar Han rats were treated with a vehicle control (Elix water) or MMV693183 by oral gavage for 7 days (Table S24) and 6 female and 6 male mice (total of 12) were used for toxicity studies, while 4 female and 4 male mice (total of 8) were used to examine toxicokinetic parameters. Body weight was measured on day 1, 4 and 8, blood was collected from the retro-orbital sinus of fasted animals under anesthesia using isoflurane in tubes containing K_3_-EDTA, citrate or Li-heparin as anticoagulant to determine the hematology, coagulation and clinical chemistry parameters, respectively, and from the jugular vein in K_2_-EDTA tubes for determining toxicokinetic parameters. Phoenix WinNonlin version 6.4 was used for Toxicokinetic evaluation.

### *In vitro* metabolism, permeability and protein binding

Stability of MMV693183 in dog, rat and hepatocytes, Caco-2 permeability, plasma protein binding and blood to plasma ratio were analyzed through a commercial service (TCG Lifesciences). The stability of MMV693183 in human hepatocytes was quantified in a relay assay (21). Briefly, cryopreserved human primary hepatocytes were thawed and cultured as described above. Hepatocytes were incubated with compounds in hepatocyte medium (0.1% DMSO) 24 h post-seeding. Supernatant was collected 1, 6 or 24 h after addition of compound, spun down and supernatant was stored at −80°C. Supernatant from the 24 h treatment was pooled and transferred to a new hepatocyte plate seeded 24 h earlier. This process was continued until the 72-h incubation time was reached. Samples were analyzed on the LC-MS/MS system Thermo Scientific™ Vanquish™ UHPLC system or Thermo Scientific™ Q Exactive™ Focus Orbitrap with a HESI-II electrospray source in positive mode using a Luna Omega Polar C18, 50 x 2.1 mm, 1.6 μm column. The chromatography was performed at a flow rate of 0.8 ml/min using a 8.70-minute gradient to 70% Mobile Phase B (0.1% formic acid in methanol) and 30% Mobile Phase A (0.1% formic acid in MilliQ water), followed by 0.40-minute gradient to 99% Mobile Phase B, and back to 1% Mobile Phase B Mobile in 0.30 min.

### Pharmacokinetic properties

The *in vivo* pharmacokinetic (PK) profiles of MMV693183 in blood and urine were determined in male CD1 mice, Sprague Dawley rats (both TCG Lifesciences) and Beagle dogs (Aptuit, Verona, Italy) (Table S24). Briefly, mice and rats were dosed intravenously or by oral gavage with MMV693183 at 3 mg/kg or 30 mg/kg, respectively. Blood sample from the saphenous vein was collected into heparinized capillary tubes at multiple intervals between 0.25 and 24 h after dosing. Subsequently, samples were spun down at 1640 g for 5-10 min at 4°C within 30 min after collection and plasma was collected. The PK study in dogs followed a cassette dosing of MMV693183, MMV693182 and MMV689258 with intravenous dosing of 1 mg/kg per compound and oral dosing of 2 mg/kg per compound. Urine was collected following an intravenous dose in both rat and dog, allowing renal clearance to be estimated. Human renal clearance was estimated from dog renal clearance as previously described (36) and other human PK parameters were predicted using allometric scaling and/or *in vitro* clearance data. PK parameters were calculated by naïve pooled approach using WinNonLin software (Phoenix, version 6.3).

### *In vivo* Pharmacokinetic-Pharmacodynamic (PKPD) relationship

The effect of MMV693183 on *P. falciparum* Pf3D7^0087/N9^ and Pf3D7^20161128TAD17N214^(48) was investigated *in vivo* using female NSG mice (Charles River) and was assessed in three different studies at TAD (The Art of Discovery, Spain) using their standard protocol (Table S24) (49). Briefly, humanized mice were engrafted daily with 0.7 ml of 50%-75% hematocrit human erythrocyte suspension until the end of the drug administration period to obtain a minimum of 40% human erythrocytes in peripheral blood during the entire experiment. Subsequently, mice were injected intravenously with 35.1×10^6^ infected RBCs in a volume of 0.3 ml. When parasitemia reached 1% (three days after infection), mice were left untreated or were treated with DHA for four days (8 mg/kg) or with MMV693183 in four dose groups: 10 mg/kg, 25 mg/kg, 50 mg/kg, and 100 mg/kg single dose by oral gavage (Table S24). When parasitemia reached the lower limit of quantitation (<0.01%) after treatment, the total circulating human RBCs were maintained by injection of human RBCs every three to four days. Parasitemia was regularly quantified (every 24 to 72 h) in each mouse by staining 2 μl of tail blood and measured on the Attune NxT Acoustic Focusing Flow Cytometer (InvitroGen) as previously described (49). The experimental designs are summarized in Table S24. Samples to quantify MMV693183 were collected and prepared at different time points depending on the study (range 0.5 to 103 h after treatment) for each mouse by mixing 25 μl of whole blood with 25 μl of Milli-Q, followed by immediate freezing of samples on dry ice. Samples were processed under liquid-liquid extraction methods and analyzed by LC-MS/MS for quantitation in a Waters UPLC-TQD (Micromass, Manchester, UK). The lower limit of quantification for MMV693183 ranged from 1 to 5 ng/ml, depending on the study.

Data preparation, exploration and model pre- and post-processing was performed using R (version 3.6.3) and R package IQRtools (version 1.2.1 IntiQuan GmbH). Non-linear mixed effects (NLME) modeling was used to estimate the PK and PD parameters using Monolix (Lixoft version MLX2018R2). The population PKPD model was developed with a two-stage approach: first a population PK model was determined; then, the individual PK parameters were used as regression parameters and the PD parameters were estimated. The PD model consists of the balance between a parasite net growth rate and a drug killing rate. The effect of MMV693183 concentration on the killing rate was estimated using an *in vitro* clearance model which is based on an E_max_ model (54) with an additional clearance term to account for the removal of dead parasites from the body and where E_max_ is fixed to the value derived from the *in vitro* parasite reduction rate (PRR) assay (Fig S16). The growth rate of log-transformed parasite concentration was fixed at 0.03 /h based on prior experiments but estimating growth rate interindividual variability. Different Hill coefficients were tested. The final model was selected based on model convergence, plausibility of parameter estimates, visual inspection of observed and model predicted time courses, standard goodness-of-fit plots and fit statistics such as Bayesian Information Criterion (BIC). The MIC and MPC_90_ were calculated with the following formulas:

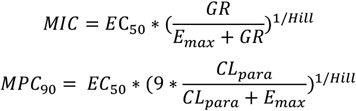

GR, parasite growth rate (1/h); MIC, minimum inhibitory concentration (ng/ml); MPC_90_, minimum parasiticidal concentration when killing/clearance rate reaches 90% of its maximum (ng/ml); EC_50_, effective concentration required to obtain 50% of maximum effect (ng/ml); Hill, Hill factor determining the steepness of the exposure-response curve; CLpara, parasitemia clearance rate (hour^-1^).

### The human efficacious dose estimation

The human efficacious dose estimation, defined as the dose able to achieve at least 9 to 12 log total parasite reduction, was predicted performing simulations using the predicted human PK parameters and the PD parameters estimated from the female NSG mice studies. Two sets of predicted human PK parameters were considered. The first set included human hepatic clearance estimated from *in vitro* hepatocyte clearance and human renal clearance estimated from dog pharmacokinetic data using allometry. The second set included the total human clearance estimated using allometry and was considered as the worst case scenario due to the higher total clearance estimated using this approach.

### Selection of drug-resistant parasites

Dd2-B2 and NF54-HGL parasites were exposed to suboptimal concentrations of MMV693183 to induce and select for drug-resistant parasites. Dd2-B2 parasites were set up at different parasite densities ranging from 10^7^-10^9^ and exposed to 3.5-9× IC_50_ (Dd2-B2 mean IC_50_ = 3.0 nM) in at least three independent experiments. Drug media was changed daily until cultures were cleared and every other day subsequently. If cultures did not clear (<0.09% parasitemia), then the concentration of drug was ramped up. Recrudescence was monitored by flow cytometry on an Accuri C6 (BD Biosciences) using SYBR Green and MitoTracker Deep Red FM (Life Technologies) as nuclear stain and vital dye, respectively. NF54-HGL parasites were treated with 3× IC_50_ for two weeks in two independent experiments and cultures were monitored by luminescence readout on the BioTek Synergy 2 Plate reader using ONE-Glo reagent (Promega). Whole-genome sequencing was performed to test for single nucleotide polymorphisms (SNPs) or copy number variations.

### Generation of transfection plasmids

The mutation found in selected drug-resistant parasites was verified by introducing the point mutations in ACS (T648M and T627A) in NF54-HGL parasites using a CRISPR-Cas9 system as described previously (14). The oligonucleotides for the guide RNA and the donor template were cloned into a pDC2-based plasmid containing a Cas9 and guide cassette, using the BbsI and EcoRI/AatII restriction sites, respectively. Donor DNA was amplified by (overlap-extension) PCR amplification from genomic *P. falciparum* DNA and oligonucleotides for the guide RNAs were ordered (Sigma-Aldrich). Correct sequence and integration of both inserts was confirmed by Sanger Sequencing.

ACS and ACS11 were C-terminally tagged with GFP using the Selection-Linked Integration (SLI) system (55). The C-terminal homology region was cloned into the plasmid using NotI/MluI restriction sites. The resulting vector was digested using EcoRV/BstZ17I restriction sites to insert the 3’ UTR. The final resulting vector contained an apicoplast-mCherry cassette that was not used in this study. Donor DNA was amplified by PCR amplification from genomic *P. falciparum* DNA. Primers are defined in Table S25.

### *Plasmodium falciparum* transfections

A DNA-loaded RBC protocol was used for transfection (56). Briefly, 100 μg of plasmid was loaded into RBCs by electroporation (310 V, 950 μF) and a trophozoite culture was added to these transfected RBCs. One day after transfection, parasites were treated with 2.5 nM WR99210 (Jacobus Pharmaceutical) for five days and cultured until they recovered. For the generation of the mutation in *ACS*, parasites were cloned by limiting dilution and integration of the mutation was confirmed by Sanger sequencing (Fig S17). For the generation of ACS- and ACS11-tagged mutants, parasites were treated with 400 μg/ml G418 (Sigma) until parasites recovered (about 15 days). Subsequently, parasites were sorted on a customized FACS Aria (BD Biosciences) based on a previously published protocol (57). In brief, a diluted sample was passed through a 70 μm nozzle. Single cells were isolated by removing doublets or cell aggregates based on FSC-H/FSC-W and SSC-H/SSC-W dot plots and selecting GFP-positive events measured using a 515 nm long pass filter. Next, 30 (ACS-tagged) or 100 (ACS11-tagged) parasites were placed back into culture. Successful integration of the transfection plasmids and absence of wild-type parasites were verified by PCR (Fig S9).

### Metabolomics assays

*P. falciparum* metabolomics analysis was performed by LC/MS as previously described (58). Briefly, parasite cultures were tightly synchronized at the ring stage one cycle prior to extraction. Trophozoite cultures at 5-10% parasitemia were purified to >90% parasitemia by magnetic purification. Parasites were counted using a hemocytometer, aliquoted to 1×10^8^ cells per condition in 5 ml medium and then placed into an incubator for 1 h to allow them to reach a metabolically stable state. Following the recovery period, MMV693183 was added at 1×, 10× or 100× IC_50_ value and compared to a no drug control in triplicate. After the incubation period, media was aspirated and the remaining culture was washed with PBS, and quenched using 90% methanol containing 0.25 μM [^13^C4^15^N]-aspartate. Blank processing samples were also quenched in the same manner to assess background metabolite levels. The samples were centrifuged, the supernatant was collected in a new tube, dried using a nitrogen gas drying rack and stored at −80 °C until they were run on the LC/MS platform.

Samples were resuspended in 1 μM chlorpropamide in 3% HPLC-grade methanol diluted in HPLC-grade water and run on a Thermo Exactive Plus Orbitrap HPLC/MS in negative mode with a scan range of 75-1000 m/z using a C18 Water Xselect HSS T3 column with 2.5 μm particle diameter. Chromatography was performed using a 25-min gradient of 3% methanol with 10 mM tributylamine and 15 mM acetic acid (solvent A) and 100% methanol (solvent B). For each analytical run, a pooled sample was generated by combining equivalent volumes of each parasite sample to assess metabolite detection and run at the beginning, middle, and end of each analytical batch to detect any possible time-dependent sensitivity changes.

### Whole genome sequencing

The Dd2-B2 parent and resistant clones were subjected to whole-genome sequencing at the Columbia University Irving Medical Center using the Illumina Nextera DNA Flex library preparation protocol and NextSeq 550 sequencing platform. Briefly, 150 ng of genomic DNA was fragmented and tagmented using bead-linked transposomes and subsequently amplified by 5 cycles of PCR to add dual index adapter sequences to the DNA fragments. The libraries were quantified, pooled and sequenced on the Illumina NextSeq high output flow cell to obtain 150 bp paired end reads.

The sequence data generated was aligned to the *P. falciparum* 3D7 genome (PlasmoDB version 36.0) using BWA (Burrow-Wheeler Alignment). PCR duplicates and reads that did not map to the reference genome were removed using Samtools and Picard. The reads were realigned around indels using Genome Analyses Tool Kit (GATK) RealignerTargetCreator and base quality scores were recalibrated using GATK Table-Recalibration. GATK HaplotypeCaller (Min Base quality score ≥ 20) was used to identify all possible variants in clones. Variants were filtered based on quality scores (variant quality as function of depth QD > 1.5, mapping quality > 30) and read depth (≥ 5) to obtain high quality SNPs that were annotated using snpEFF. The list of variants from the resistant clones were compared against the Dd2-B2 parent to obtain homozygous SNPs present exclusively in the resistant clones. Copy number variations were detected using the BicSeq package by comparing the read counts of the resistant clones against the Dd2-B2 parent. Integrative Genomics Viewer was used to verify the SNPs and copy number segments in the resistant clones.

Genomic DNA from the parental NF54 line and the MMV693183-induced resistant lines were sequenced at the Pennsylvania State University according the Illumina® Truseq Sequencing protocol. Following sequencing, the data were processed using the Tadpole Galaxy scientific data analysis platform (59). Briefly, the Trimmomatic tool was used to trim adapter sequences and the genome was mapped using the Map with BWA-MEM tool against a *P. falciparum* 3D7 reference genome. The Filter Sam or Bam, output Sam or Bam tool was used to consolidate the reads and generate the BAM file for the remaining analyses. The Depth of Coverage on Bam File tool was used to assess the depth of coverage of the dataset. Finally, the Freebayes – Bayesian genetic variant detector tool was used to assess the data for SNPs, inserts, and deletions.

### Generation of antigen for ACS antibody production

Recombinant protein fragment used for immunization was obtained by cloning the first 414 nucleotides of a codon-optimized coding sequence of ACS in frame with an N-terminal Glutathione S-transferase (GST)tag, into the expression vector pGex-4T. The vector was transformed into competent *Escherichia coli* BL21 (DE3) expression cells to express a recombinant protein fragment. Protein production was induced in 250 ml log-phase growing cells with 500 μM IPTG for 3 h at 30°C. After incubation, the cells were collected, resuspended in 10 ml 20 mM Tris-HCl (pH 7.5) and disrupted by sonicating 3 times for 45 seconds on ice. The protein was released from the cell debris using 8 M urea and dialyzed in 10 mM Tris-HCl (pH8.1) with 0.1% Triton X-100. The samples were stored at −20 °C prior to immunization.

### Generation of polyclonal antiserum and immunoprecipitation assays

Rabbits were immunized with recombinant ACS according to the manufacturer’s standard procedures (Eurogentec, Seraing, Belgium). Reactivity of serum was compared to pre-immune serum using an enzyme-linked immunosorbent assay. Briefly, plates were coated with 100 ng antigen per well and a dilution range of serum (pre-immune versus serum from final bleed) was added. Antibody binding was measured with a biotinylated goat anti-rabbit secondary antibody using the Vectastain ABC kit (Vector Labs). Immunoglobulins were absorbed on protein A/G sepharose (Pierce) and used to isolate ACS from *P. falciparum* parasite lysates.

### Parasite lysates for ACS assay

Asynchronous blood-stage *P. falciparum* strain NF54 was released from the RBCs by incubation with 0.06% saponin in PBS for 5 min on ice. Parasites were pelleted by centrifugation (10 min at 4,000xg), washed with PBS and lysed in 50 mM NaF, 20 mM Tris-HCl (pH 7.5), 0.1% Triton X-100, 2 mM dithiothreitol, 2 mM EDTA and 1% (v/v) Halt Protease Inhibitor Cocktail (Thermo-Fischer Scientific, Waltham, MA, USA). Suspensions were then sonicated 6 times for 3 seconds at an amplitude of 16 microns peak-to-peak. Sonicated samples were centrifuged at 24,000 g for 5 min at 4°C and supernatants were used in immunoprecipitation and enzyme activity assays.

### ACS assay

Acetyl-CoA synthetase activity was measured using a radioactively labeled ACS assay. The reaction mixtures contained 8 mM MgCl_2_, 2 mM ATP, 30 μM Coenzyme A, 200 μM ^14^C-labeled sodium acetate (PerkinElmer), 50 mM Hepes-KOH, pH 8.5 and immunoprecipitated ACS in a total volume of 35 μl. Reactions were incubated at 37°C for 30 min. The reaction was terminated with 3.5 μl of a 10% acetic acid solution in 90% ethanol. Samples were loaded on DEAE filter paper (GE Healthcare) and washed thoroughly in 2% acetic acid solution in 95% ethanol to wash away unreacted acetate. After the discs were dried, they were transferred into scintillation vials containing 3 ml ScintiSafe 30% Cocktail (Fischer Scientific, Hampton, NH, USA). Radioactivity in each vial was counted using a Tri-Carb 2900TR Liquid Scintillation Analyzer (Packard Bioscience, Boston, MA). To test the inhibitory properties of MMV693183, 4’P-MMV693183, and CoA-MMV693183 on ACS, a dilution range of the compound was pre-incubated for 30 min with the immunoprecipitated ACS before initiation the ACS reaction by adding the reaction mixture. The 4’P-MMV693183 and CoA-MMV693183 metabolites were obtained from Syncom, Groningen, the Netherlands.

### Cellular Thermal Shift Assay

A cellular thermal shift assay (CETSA) was performed on infected RBCs or parasite lysates as described previously (26).

For CETSA on infected RBCs, synchronized trophozoite cultures (NF54-HGL parasites) were purified using magnetic-activated cell sorting (MACS). Parasites were resuspended in 1× PBS, aliquoted in PCR tubes (1.8×10^7^ cells/tube) and subjected to 37°C or 51°C for 3 min on a pre-heated PCR machine, followed by 4°C for 3 min. Parasites were mixed with 2× lysis buffer (100 mM HEPES, 10 mM β-glycerophosphate, 0.2 mM activated Na3VO4, 20 mM MgCl2, with EDTA-free protease inhibitor cocktail (Merck)), and subjected to three freeze-thaw cycles using liquid nitrogen, followed by mechanical shearing using a syringe with a 25G needle. Samples were spun down at 18,000 g for 20 min at 4°C and the soluble fraction was flash-frozen in liquid nitrogen and stored at -−80°C.

For a CETSA on parasite lysates, synchronized trophozoite cultures (ACS11-tagged parasites) were treated with 0.1% saponin to lyse the RBCs, and washed three times in PBS. Subsequently, parasite pellets were resuspended in 1× lysis buffer and subjected to three flash-freeze-thaw cycles using liquid nitrogen, followed by mechanical shearing using a syringe with a 25G and a 30G needle. Samples were spun down at 18,000 g for 20 min at 4°C. Supernatant was diluted to 2.1 mg/ml protein concentration, and 100 μl was added to each PCR tube containing 1 μl of compound at a 100× concentration (final concentration of 1 μM). Samples were incubated for 30 min at room temperature, subjected to a thermal gradient for 3 min on a pre-heated PCR machine, followed by 4°C for 3 min. Samples were spun down at 18,000 g for 20 min at 4°C and the soluble fraction was flash-frozen in liquid nitrogen and stored at −80°C.

### Western blot

Samples were loaded on an 8% SDS-Page gel (Genscript) with 4×10^6^ infected RBCs or 50 μg protein for the parasite lysate approach. Proteins were transferred to a nitrocellulose membrane that was blocked with 5% skim milk (Sigma) in PBS overnight and incubated with rabbit antiserum against ACS (1:1000) for 1 h. Subsequently, membranes were washed three times with PBS-Tween for 5 min, followed by incubation with secondary horseradish peroxide (HRP)-conjugated goat anti-rabbit antibodies (Dako P0448, 1:1000) for 1 h. Blots were then washed three times with PBS-Tween for 5 min and twice with PBS. Following a 5-min incubation with Clarity Max Western ECL Substrate (BioRad), protein blots were imaged using the ImageQuant LAS4000 (GE Healthcare). The band intensity was quantified using Fiji software.

### Immunofluorescence microscopy

Asynchronous asexual blood-stage ACS-GFP or wild-type NF54 parasites were allowed to settle on poly-L-lysine coated coverslips for 20 min at room temperature. Parasites were fixed with 4% EM-grade paraformaldehyde and 0.0075% EM-grade glutaraldehyde in PBS for 20 min and permeabilized with 0.1% Triton X-100 for 10 min (60). Samples were blocked with 3% bovine serum albumin (BSA) (Sigma-Aldrich) in PBS for 1 h. Samples of ACS-GFP and NF54 parasites were incubated with primary chicken anti-GFP antibody (1:100, Invitrogen), or ACS pre-immune and immune serum (1:500), respectively, in 3% BSA/PBS for 1 h, followed by incubation with secondary goat anti-chicken AlexaFluor 488 antibody (1:200, Invitrogen) in 3% BSA/PBS for 1 h. Nuclei were visualized with 1 μM DAPI in PBS for 1 h. PBS washes were performed between different steps. Coverslips were mounted with Vectashield (Vector Laboratories). Images were taken with a Zeiss LSM880 Airyscan microscope with 63x oil objective with 405 and 488 nm excitations. Images were Airyscan processed before analysis with FIJI software. Since no quantitative comparisons were performed, brightness and contrast were slightly altered in FIJI to improve visualization of AlexaFluor488 and DAPI signals.

### Statistics

Dose-response assays were analyzed by a nonlinear regression using a four-parameter model and the least squares method to find the best fit. One-way Analysis of variance (ANOVA) was performed using the Bonferroni’s Multiple Comparison Test.

## Acknowledgements

We gratefully acknowledge A. Fuchs for the PKPD analyses of pantothenamides that guided the selection of the preclinical candidate, S. Mok for assistance with whole-genome sequencing analysis, C. Bioni for providing access to the laboratory for the Brazilian-field isolates *ex vivo* assessments, O. Byaruhanga, S. Orena, M. Okitwi and T. Katairo for assistance with *ex vivo* assays on fresh *P. falciparum* isolates in Uganda, S. Sax for technical assistance with the SCID mouse *P. falciparum in vivo* efficacy studies performed at Swiss TPH. We thank D.F. Wirth, A.K. Lukens and R. Summers for pre-publication sharing of data and fruitful discussions. T. Spielmann is acknowledged for providing the plasmid SLI-TGD. We also thank the Huck Institutes of Life Sciences Metabolomics Core Facility at Penn State University.

## Author contributions

T.W.A.K, and K.J.D. conceived the work and did overall supervision and analysis of parasitology, biochemistry, and molecular biology. L.E.d.V. performed and analyzed molecular biology experiments and *in vitro* parasitology assays and generated parasite mutants, T.W.A.K. provided supervision. J.M.J.V. generated parasite mutants and performed and analyzed immunofluorescence assays, L.E.d.V. and T.W.A.K provided supervision. P.A.M.J. performed and analyzed molecular biology and biochemistry assays, J.S. provided supervision. C.B. generated and analyzed pharmacokinetic - pharmacodynamic models. J.M. performed and analyzed metabolomics data and whole genome sequencing data, M.L. provided supervision. S.W., M.B.J.-D., and I.A.-B. performed and analyzed efficacy and pharmacokinetics studies in SCID mice. C.F.A.P performed and analyzed growth assays on knockdown parasite mutants, J.C.N. provided supervision. J.M.B., R.H., T.H. K.M.J.K. performed and analyzed *in vitro* parasitology assays, K.J.D. provided supervision. K.R., J.S., T.Y. generated resistant parasite lines, performed and analyzed *in vitro* parasitology experiments and whole genome sequencing data, D.A.F. provided supervision. G.T. designed and analyzed experiments to predict human PK parameters. B.C.F. performed and analyzed the parasite reduction rate assay, L.M.S., F.J.G. provided supervision. A.C. performed and analyzed the dual gamete formation assay, J.B. provided supervision. R.R. designed and analyzed the hemolytic toxicity assay. A.C.C.A., D.B.P., P.K.T. performed and analyzed *ex vivo* parasitology assays, R.V.C.G. R.A.C., P.J.R. provided supervision. P.H.H. overall supervised medicinal chemistry. R.B., B.C., R.W.S., and J.S. advised on parasitology and drug development. L.E.d.V., T.W.A.K. and K.J.D. wrote the manuscript. All authors proofread and edited the manuscript.

