## SupplementaryInformation for "Preclinical characterization and target validation of the antimalarial pantothenamide MMV693183"

[1]Department of Medical Microbiology, Radboudumc Center for Infectious Diseases, Radboud Institute for Molecular Life Sciences, Radboud University Medical Center, Nijmegen, The Netherlands. [2]Department of Dermatology, Radboud Institute for Molecular Life Sciences, Radboud University Medical Center, Nijmegen, The Netherlands. [3]Medicines for Malaria Venture, Geneva, Switzerland. [4]Department of Chemistry and Huck Center for Malaria Research, The Pennsylvania State University, University Park, PA 16802, United States of America. [5]Department of Biological Engineering, Massachusetts Institute of Technology, 77 Massachusetts Avenue, Cambridge, MA 02139, United States of America. [6]Department of Microbiology & Immunology, Department of Medicine, Columbia University Irving Medical Center, New York, New York. [7]TropIQ Health Sciences, Nijmegen, The Netherlands. [8]Infectious Diseases Research Collaboration, Kampala, Uganda. [9]Sao Carlos Institute of Physics, University of São Paulo, São Carlos, São Paulo, Brazil, Av. João Dagnone, 1100, 13563-120 São Carlos-SP, Brazil. [10]The Art of Discovery, Derio, Spain. [11]Department of Life Sciences, Imperial College London, South Kensington, London, SW7 2AZ, UK. [12]Global Health, GlaxoSmithKline, Tres Cantos, Madrid, Spain. [13]Research Center for Tropical Medicine of Rondonia, Porto Velho, Brazil, Av. Guaporé, 215, Porto Velho- RO, 76812-329, Brazil. [14]Department of Immunology and Microbiology, University of Colorado Anschutz School of Medicine, United States of America. [15]Sygnature Discovery, Nottingham, United Kingdom. [16]Swiss Tropical and Public Health Institute, Basel, Switzerland. [17]University of Basel, Basel, Switzerland. [18]Department of Natural Sciences and Mathematics, Dominican University of California, San Rafael, CA, USA. [19]Department of Medicine, University of California, San Francisco, CA, USA. [20]Hermkens Pharma Consultancy, Oss, The Netherlands. [21]Department of Biochemistry & Molecular Biology, The Pennsylvania State University, University Park, PA 16802, United States of America. *Contributed equally, Corresponding authors.

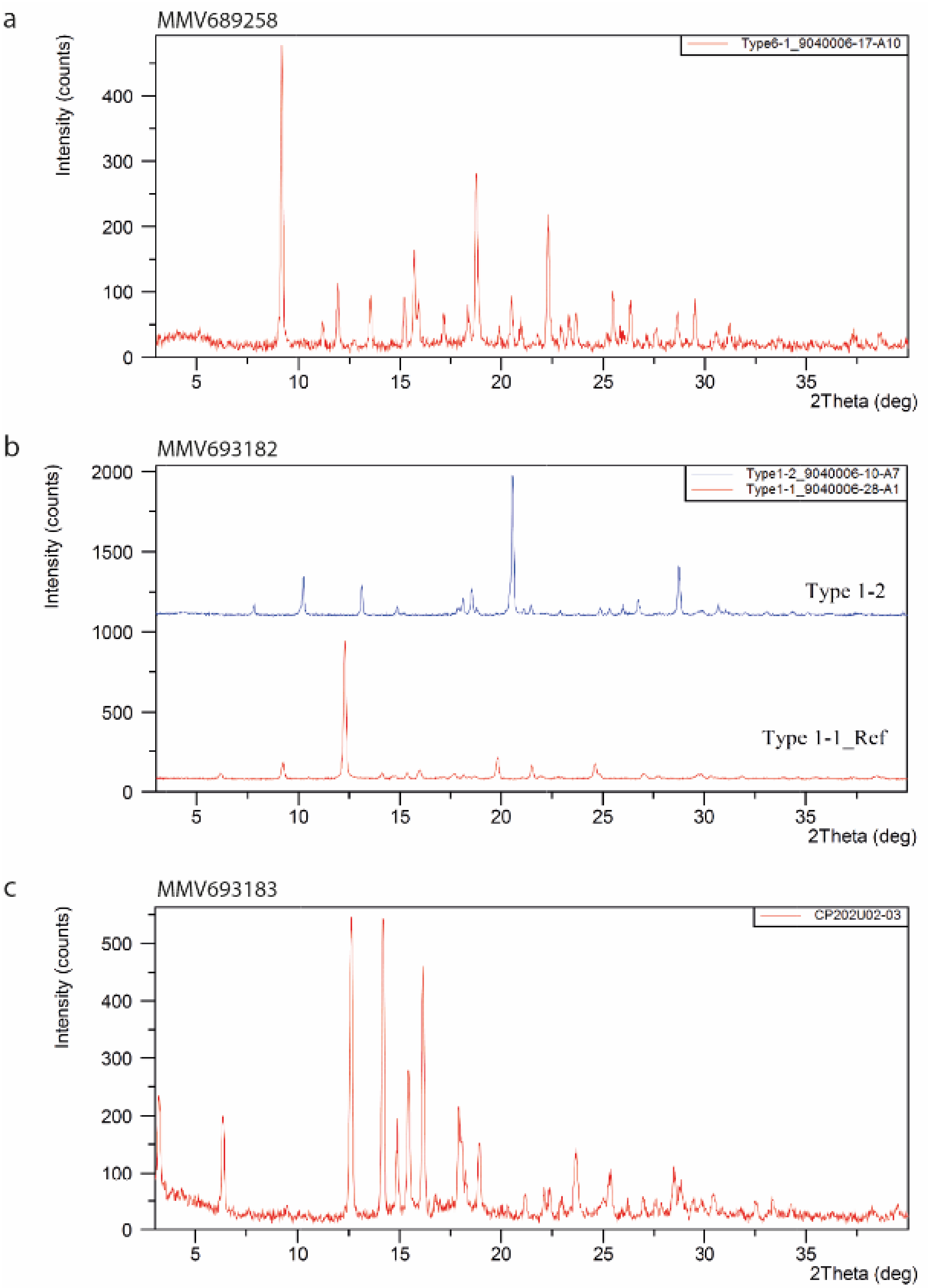

**Fig S1. XRPD patterns from crystallization screen for crystalline MMV689258 (a), MMV693182 (b) and MMV693183 (c).**

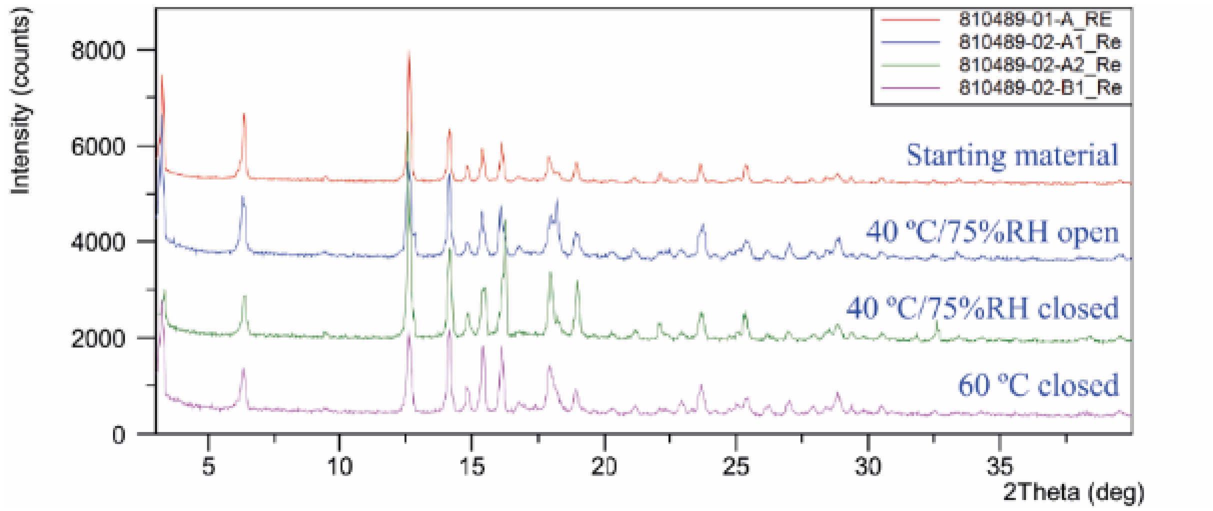

**Fig S2. XRPD overlay of stability samples.** MMV693183 was incubated at different conditions of stress for one month. XRPD patterns show that there is no form-change from the starting material, indicating the stability of this crystalline product.

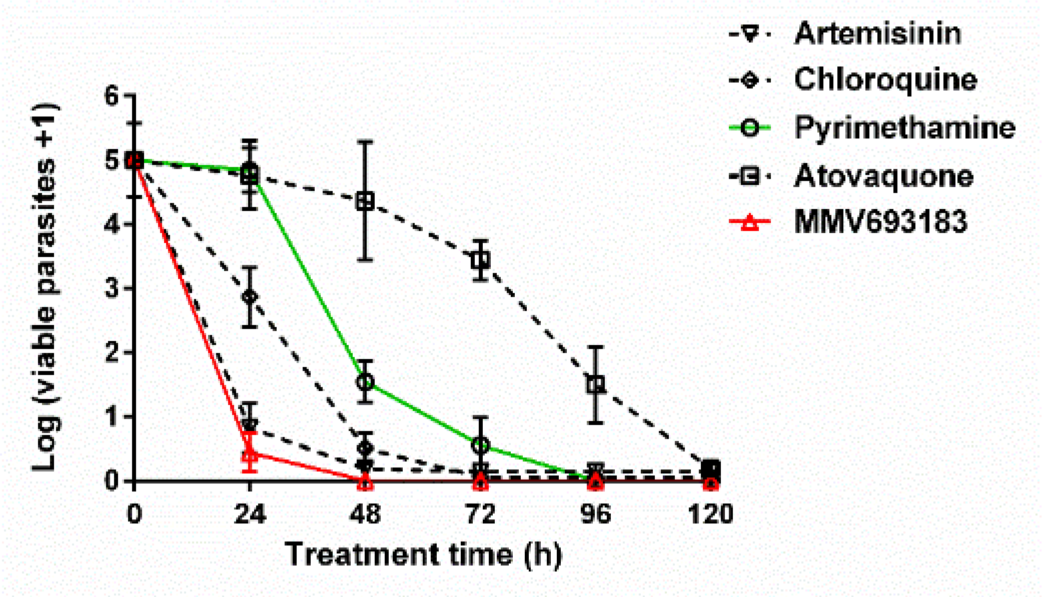

**Fig S3. *In vitro* antimalarial killing rate.** Parasites were treated with 40 nM MMV693183 or 0.94 µM pyrimethamine and the number of viable parasites were quantified in one experiment with four technical replicates (±SD). Artemisinin, chloroquine and atovaquone are retrieved from different datasets and shown for comparative purposes.

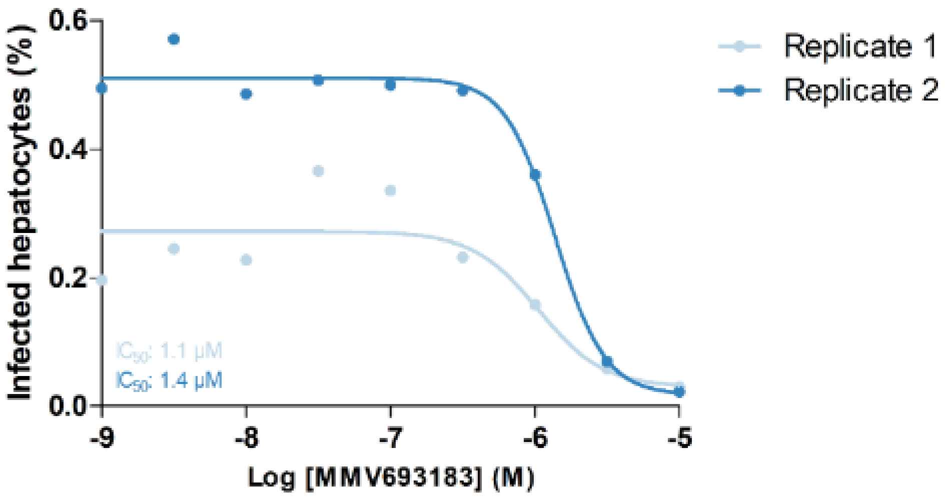

**Fig S4. MMV693183 is not efficacious against liver stages.** Cryopreserved human primary hepatocytes were infected with *P. falciparum* sporozoites and exposed to different concentrations of MMV693183. The number of infected hepatocytes were quantified using microscopy. Data from one experiment with two replicates are shown. While a difference in infected hepatocytes was observed within 1 experiment, both replicates are shown. Reassuringly, both replicates have a similar $IC_{50}$ value.

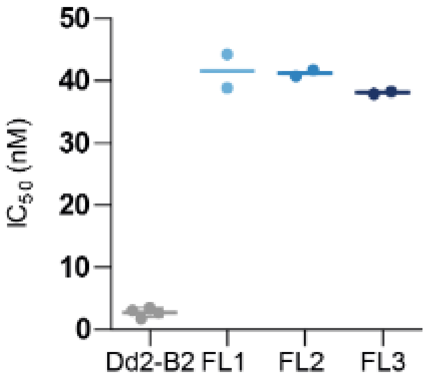

**Fig S5. MMV693183 resistant bulk cultures in three independent cultures**. The mean $IC_{50}$ (±SEM) of Dd2-B2 parasites and three independent MMV693183-resistant bulk cultures (Fl1, Fl2, Fl3) against MMV693183 is determined from two to four independent experiments measured in technical duplicates.

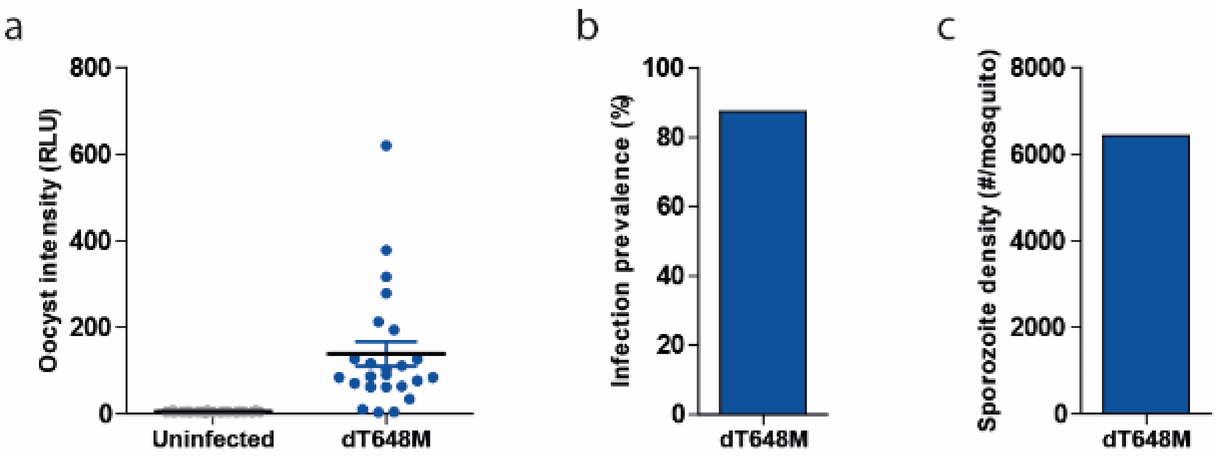

**Fig S6. Transmission of MMV693183-resistant *P. falciparum*.** Mosquito infections of the drug-induced resistant mutant harboring a mutation in ACS (T648M). The oocyst intensity is depicted as relative luminescent units (RLU) (**a**), infection prevalence (n = 1) (**b**) and sporozoite density (n=1) (**c**) after mosquito feeds.

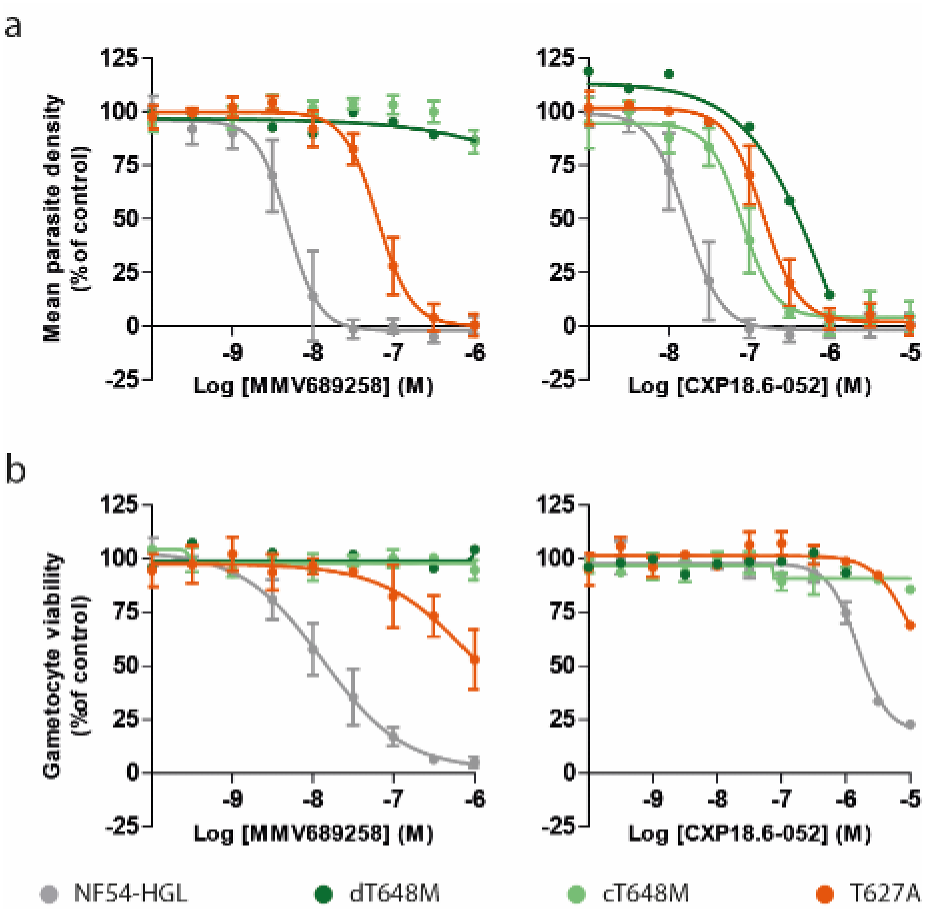

**Fig S7. Cross-resistance of ACS mutants to pantothenamides**. (**a-b**) Drug sensitivity assays are performed on asexual (**a**) or sexual blood-stage parasites (**b**) with a T648M or T627A mutation in ACS. The MMV693183-induced resistant parasite line (dT648M) was exposed to MMV689258 and CXP18.6-052 in one experiment with two technical replicates and the CRISPR-engineered parasites (cT648M and T627A) were exposed to MMV689258 or CXP18.6-052 in three independent experiments (two technical replicates per experiment). The average value to control ± SEM is presented.

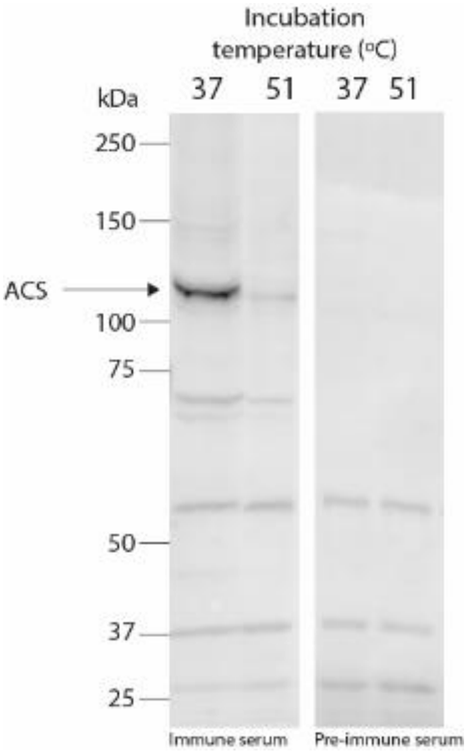

**Fig S8. Reactivity of immune serum used to extract ACS from parasite lysate.** Western blot of NF54-HGL parasites incubated with rabbit pre-immune or immune sera directed against a recombinant ACS fragment. Parasites were MACS-purified and incubated for 3 min at 37°C or 51°C, followed by incubation at 4°C for both conditions. Subsequently, parasites were lysed using lysis buffer for CETSA assays, three freeze-thaw cycles, and trituration through a 25G needle. About $4\times10^6$ cells were loaded per lane and the blot was either incubated with immune serum (left) or pre-immune serum (right). The intensity of the non-specific bands is an indicator of equal loading. ACS is indicated by the arrow and is predicted to be ~113.8 kDa.

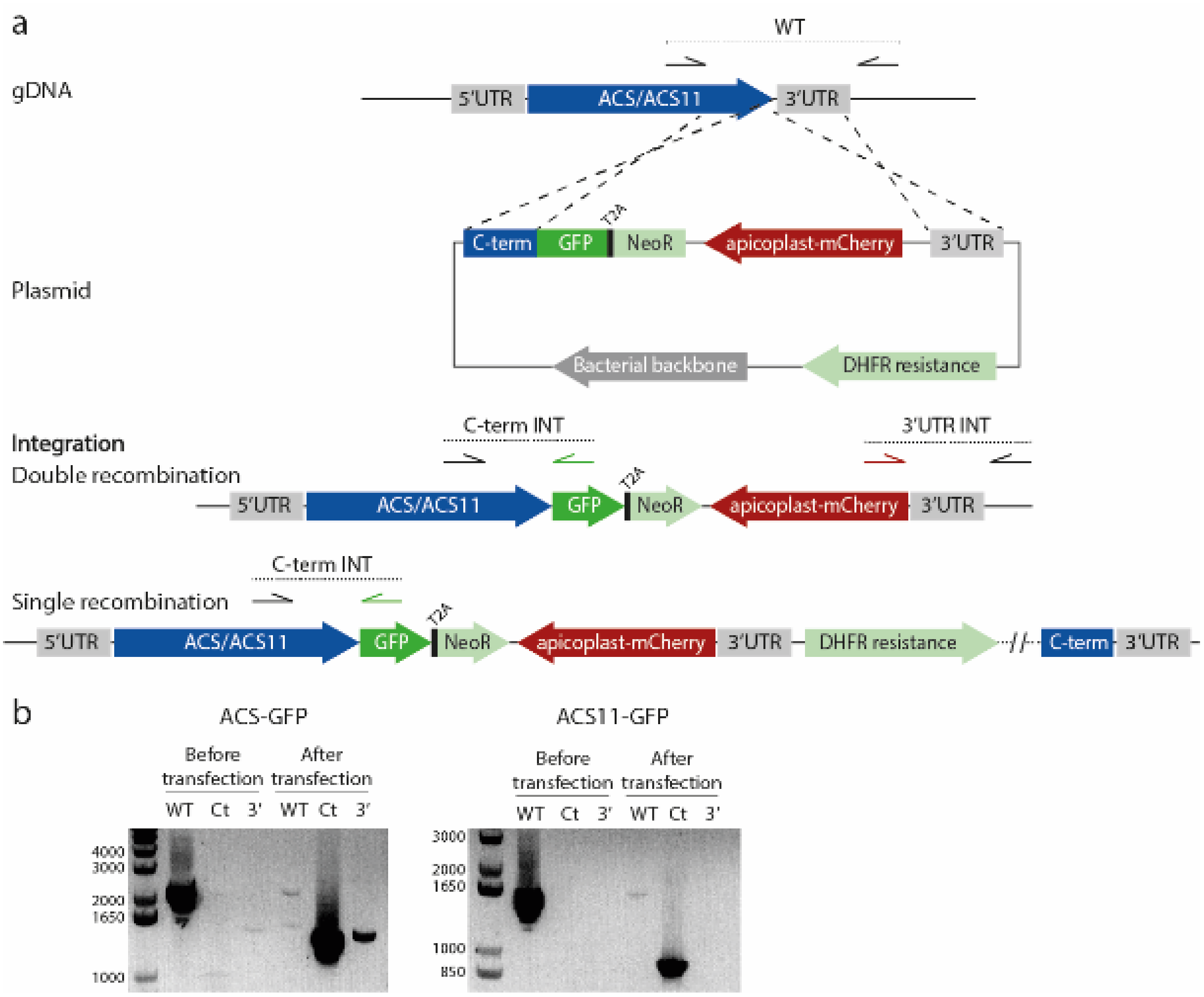

**Fig S9. Generation of ACS and ACS11 fusion with GFP.** (**a**) Schematic of the GFP-tagging approach of *ACS* and *ACS11* using selection linked integration. There are two possibilities for integration, double and single recombination, and PCR products are indicated for wild-type or integration. (**b**) Diagnostic PCR confirmed that *ACS* and *ACS11* are both fused to GFP. The plasmid is integrated in *ACS11* only by single recombination, while it is integrated by double recombination, and possibly single recombination, in *ACS*.

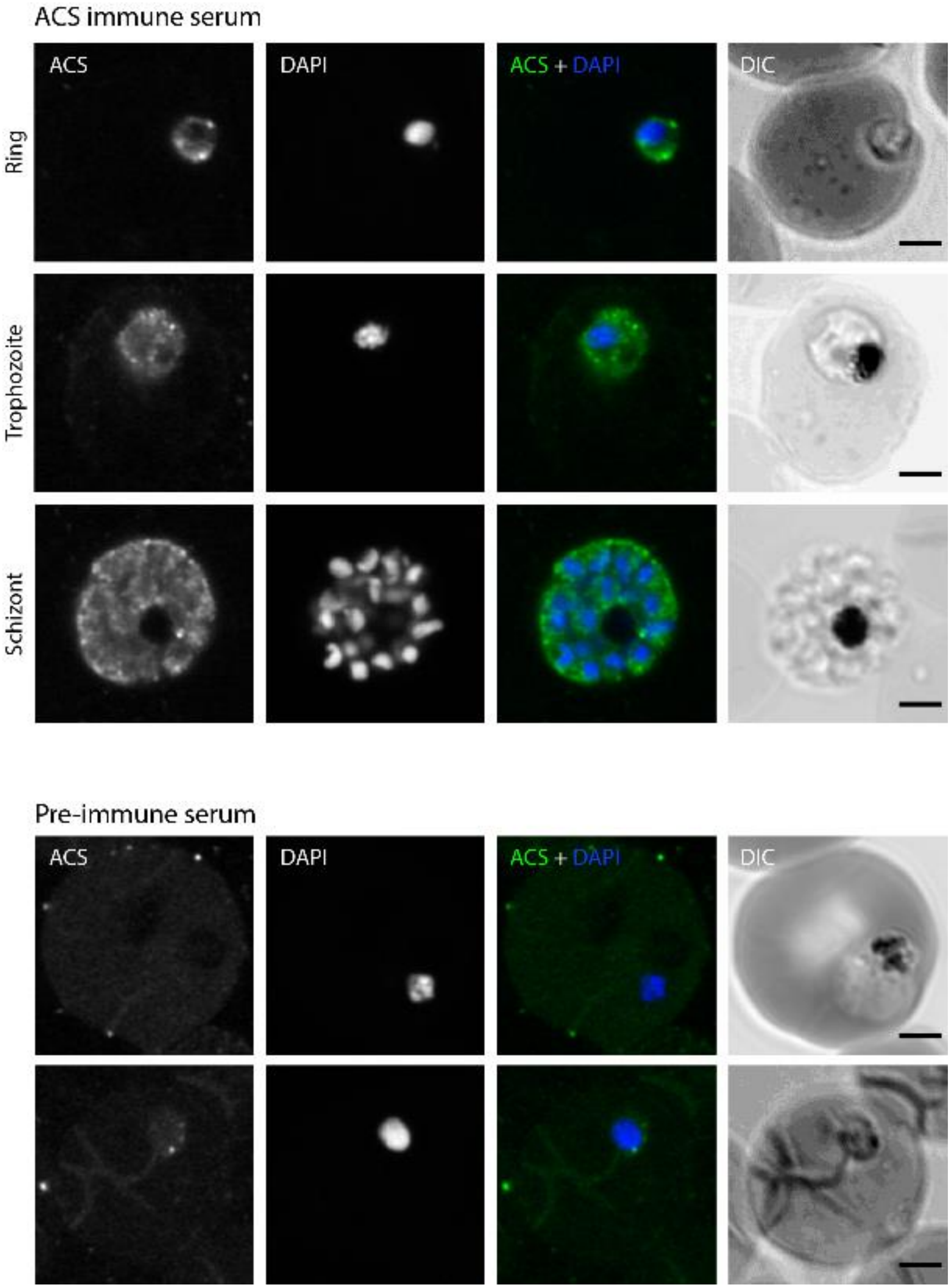

**Fig S10. Immunofluorescence microscopy of wild-type *Plasmodium falciparum* strain NF54 asexual blood-stage parasites stained with ACS pre-immune or immune serum.** Depicted are representative images of parasites stained with rabbit ACS immune serum (top) or pre-immune serum (bottom). DNA was stained with DAPI. Scale bars, 2 µm.

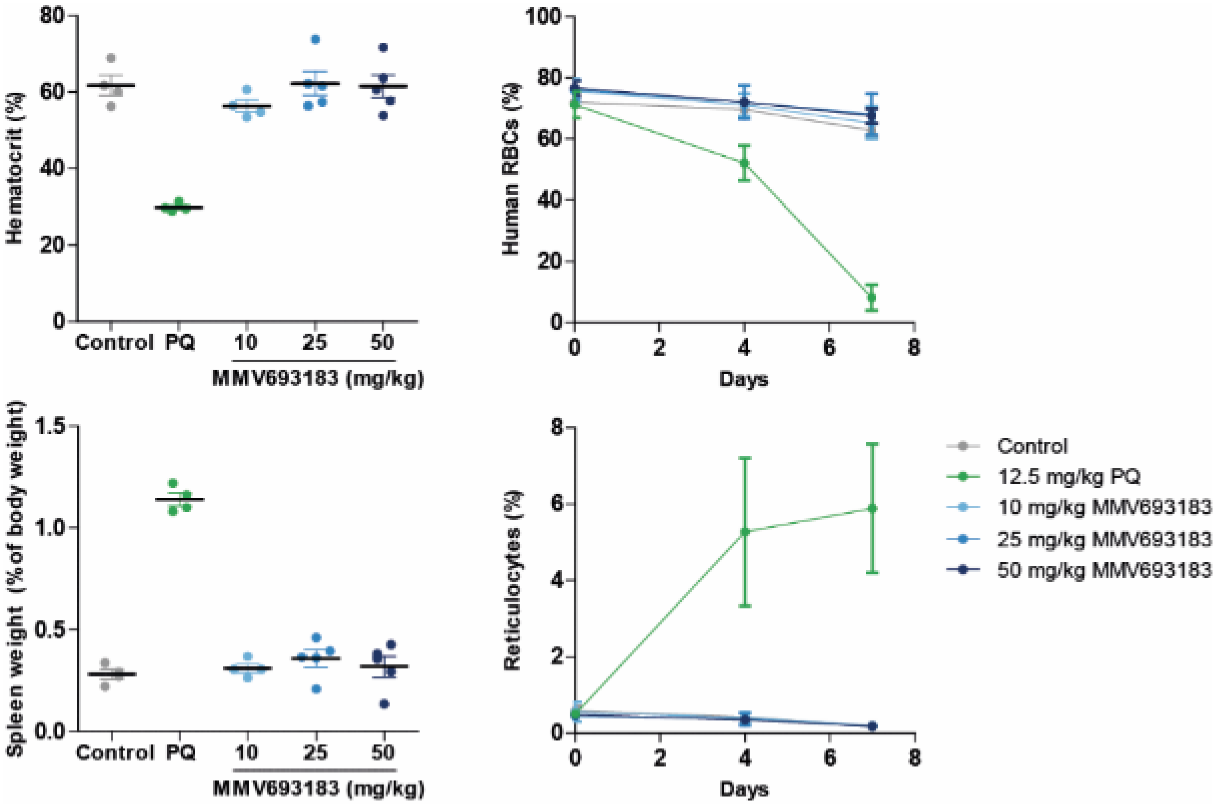

**Fig S11. Absence of detectable hemolytic toxicity in G6PD-deficiency mouse model**. The hemolytic activity of MMV693183 was determined in NSG mice engrafted with human RBCs from an A- G6PD-deficient donor with primaquine as a positive control. The percentage hematocrit, human RBCs and reticulocytes, and the spleen weight normalized to body weight ±SD are shown for N = 4-5 mice/treatment.

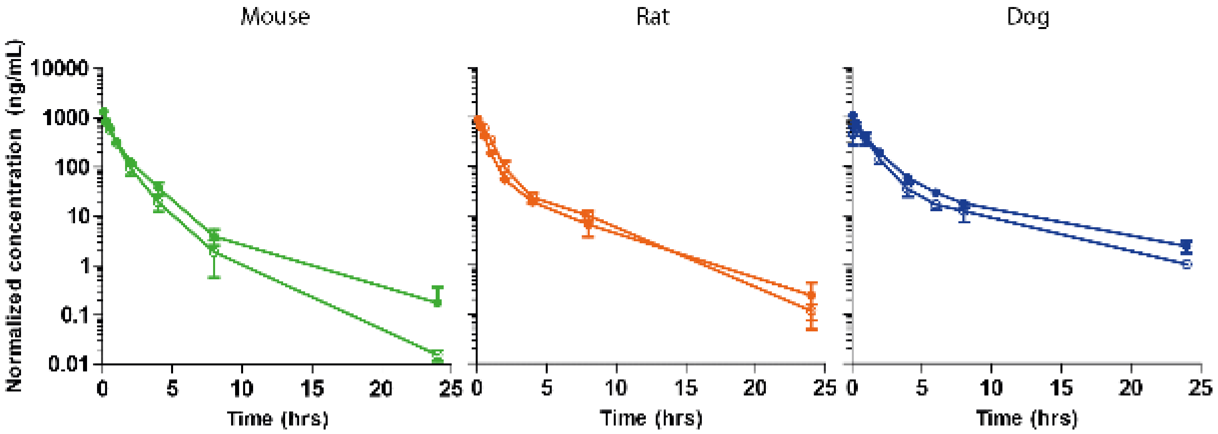

**Fig S12. Pharmacokinetics in mice, rats and dogs.** The plasma concentration (±SD) of MMV693183 in mice, rats and dogs was determined and is presented as the concentration of compound normalized to the dose received for three animals per treatment. Filled circles represent intravenous injection (3 mg/kg for mice and rats, 1 mg/kg for dogs) and clear circles represent *per os* dosing (30 mg/kg for mice and rats, 2 mg/kg for dogs).

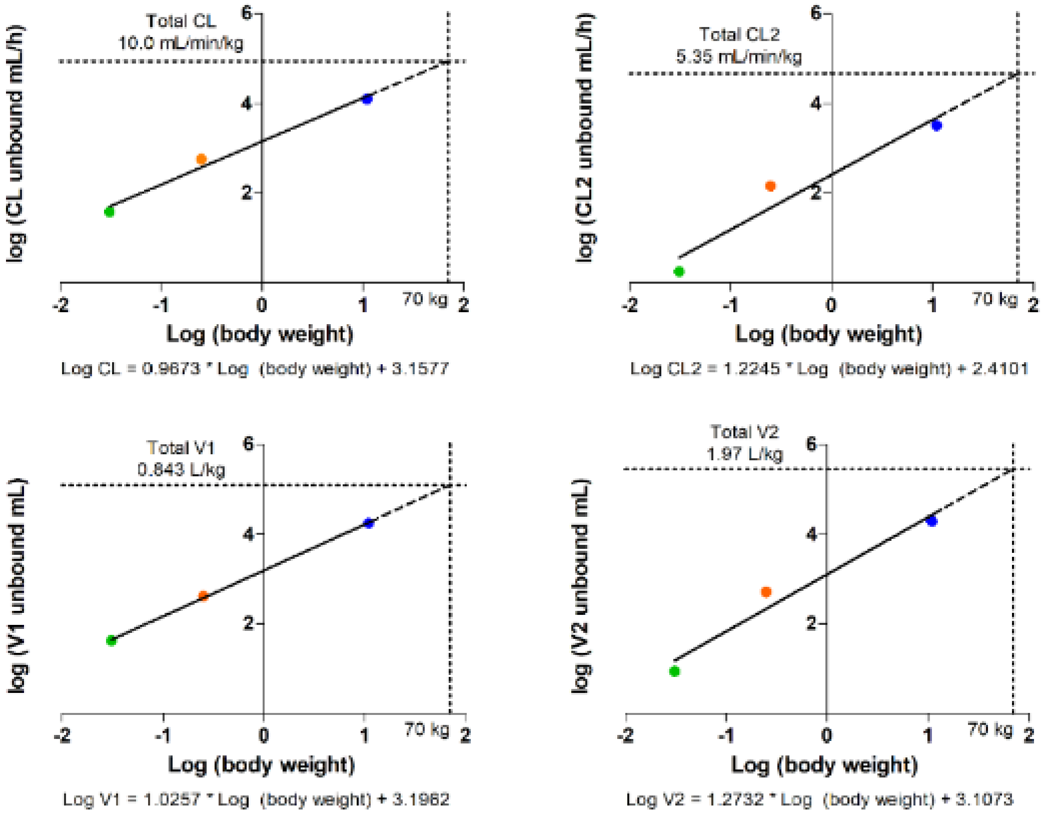

**Fig S13. Prediction of human pharmacokinetic values using allometry scaling.** The average unbound plasma concentration was calculated in Phoenix 64 using a two-compartment model with multiplicative weighting parameterized by clearance and volume of distribution in the central (CL; V1) or effect (CL2; V2) compartment in mice (green), rats (orange) and dogs (blue). Allometry scaling is used to predict the human parameters.

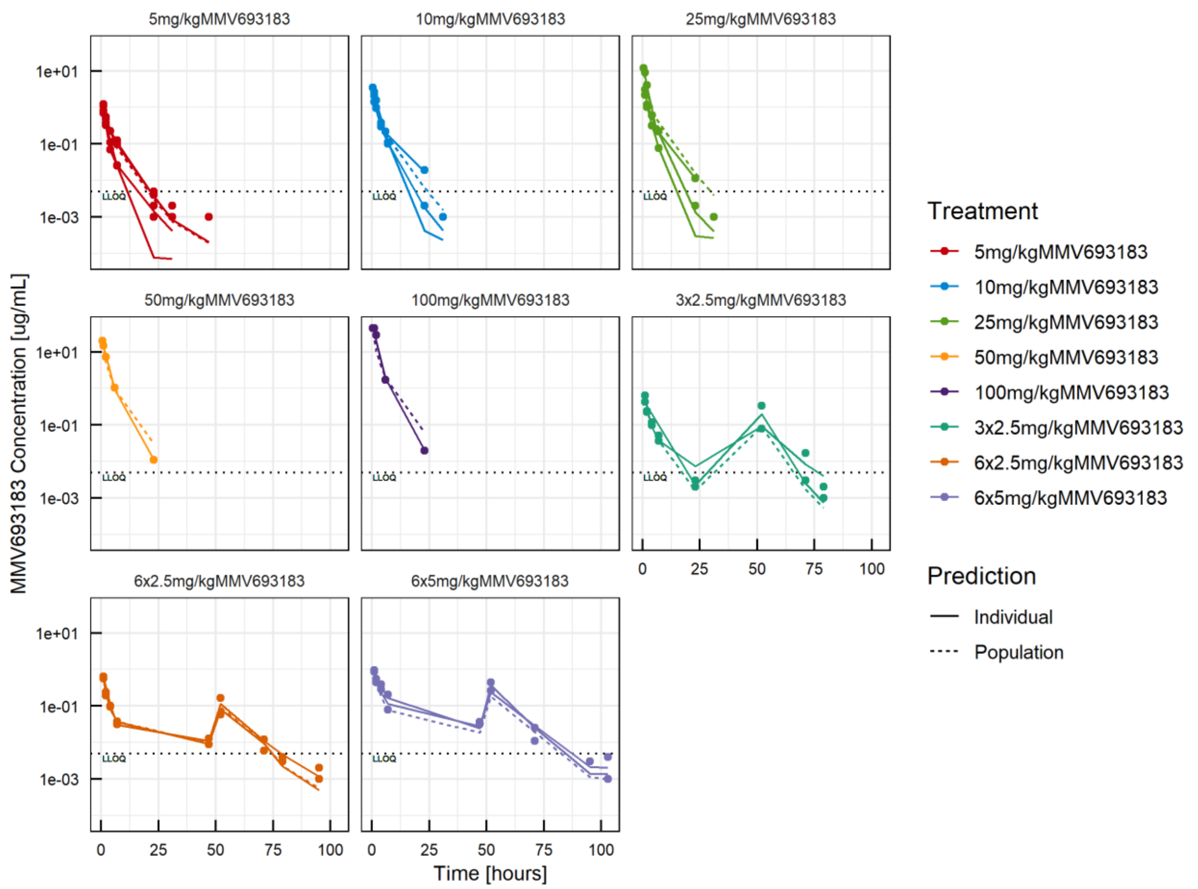

**Fig S14. PK individual fits from humanized mice.** PK data in NSG mice were calculated using an NLME approach. A 3-compartment model with zero-order absorption, linear elimination and a combined residual unexplained variability was selected as the best PK model describing the PK data. There was no lag time observed for drug absorption. Observations are represented by full circles, continuous lines represent individual predictions, while dashed lines represent the PK model at a population level.

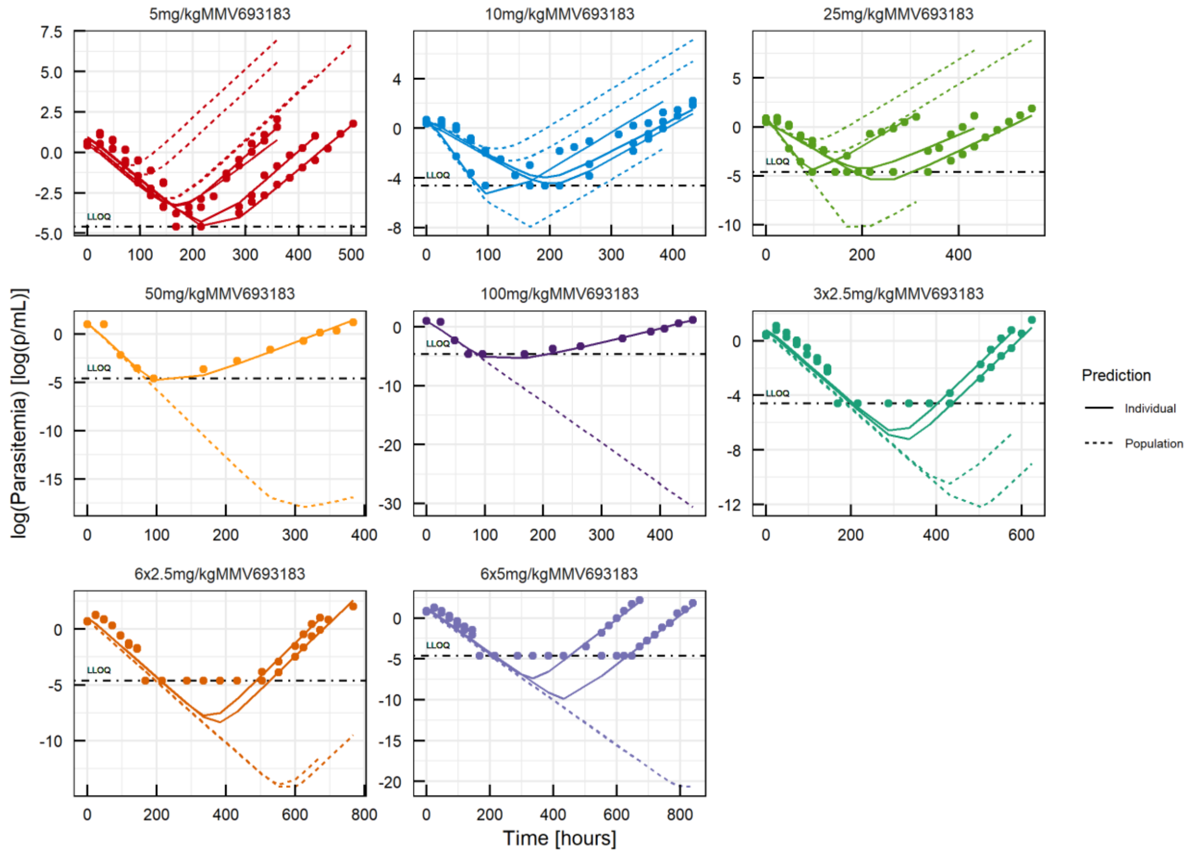

**Fig S15. PD individual fits using the *in vitro* clearance model.** Observations are represented by full circles, continuous lines represent individual predictions, while dashed lines represent the PK model at a population level.

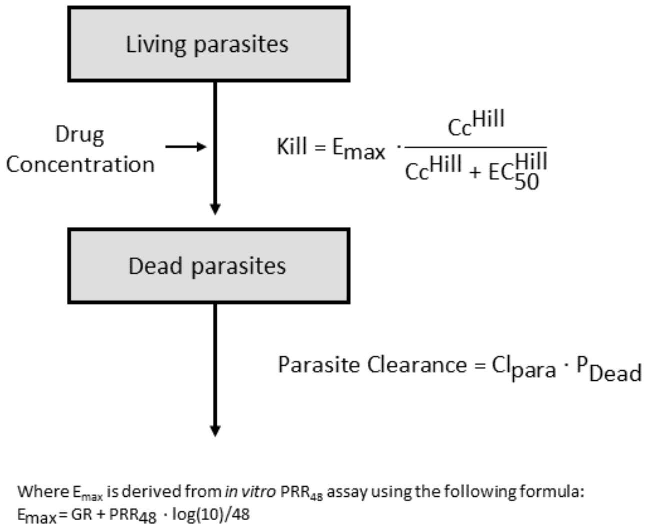

**Figure S16.** Description of the *in vitro* clearance model used to fit the NSG mice data

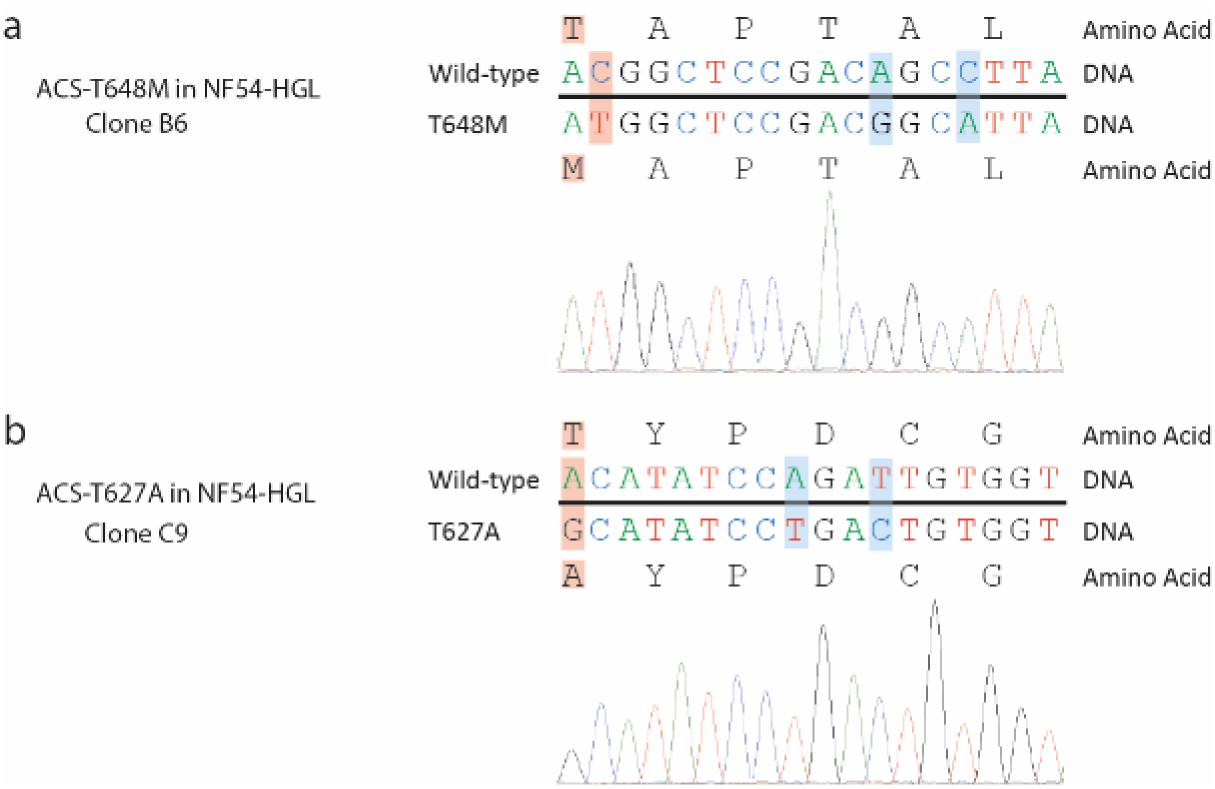

**Fig S17. Sequence verification of CRISPR-Cas9 engineered mutants.** The *ACS*-T648M (**a**) and the *ACS*-T627A (**b**) mutation were introduced in NF54-HGL parasites and the integration was confirmed by Sanger sequencing.

**Table S1. PK parameters derived from the data shown in Fig 1**

| Compound | $T_{max}$ (h) | $C_{max}$ (ng/ml) | $AUC_{0-24h}$ (h*ng/ml) |
|---|---|---|---|
| MMV689258 | 1.00 | 3780 | 6730 |
| MMV693183 | 1.50 | 12900 | 33700 |
| MMV884962 | 1.00 | 3920 | 8650 |
| MMV693182 | 1.00 | 5270 | 15100 |
| MMV976394 | 1.00 | 5810 | 10800 |
| MMV1542001 | 1.00 | 5150 | 11800 |

**Table S2. Solubility of MMV693183**

| Solution | Solubility (ppm) |
|---|---|
| **PBS** | 7137.52 |
| **FaSSIF** | 9248.25 |
| **FeSSIF** | 9050.89 |

**Table S3. Selection of MMV693183-resistant parasites in *P. falciparum*.**

| Inoculum and strain | Concentration MMV693183 for selection | Frequency (# of parasite positive flasks or wells) | $IC_{50}$ shift | Mutation |
|---|---|---|---|---|
| $1x10^5$ Dd2 | 15 nM | 0/24 wells | N/A | N/A |
| $1x10^7$ Dd2 | 15-22 nM (1x)<br>10.5-14 nM (2x) | 0/3 flasks | N/A | N/A |
| $1x10^8$ Dd2 | 10.5-1 4nM | 0/3 flasks | N/A | N/A |
| $1x10^9$ Dd2 | 12-27 nM | 3/3 flasks | 13x | Single SNP T648M in PF3D7_0627800 (ACS) (3/3) |
| NF54 | 7 nM | 2/2 flasks | 77x | T648M in PF3D7_0627800 (ACS) (2/2) |

**Table S4. Mutations in MMV693183-resistant NF54 parasites**

| Chromosome | Position | GeneID | Annotation | Mutation |
|---|---|---|---|---|
| PF3D7_04_v3 | 374669 | PF3D7_0407600 | Conserved *Plasmodium* protein, unknown function | Non_Synonymous_Coding (ATA->ACA I541T; I544T) |
| PF3D7_05_v3 | 1332563 | | | NonCoding - upstream |
| PF3D7_06_v3 | 52711 | PF3D7_0601400 | Erythrocyte membrane protein 1 (PfEMP1), pseudogene | Stop_Gained (TTG->TAG L454*) |
| PF3D7_06_v3 | 1115595 | PF3D7_0627800 | Acetyl-CoA synthetase | Non-Synonymous_Coding (ACT->ATG T648M) |
| PF3D7_09_v3 | 805836 | PF3D7_0919800 | TLD domain-containing protein | Codon_Change_Plus_Codon_Insertion TTATTA insert disrupting AA 606 |
| PF3D7_14_v3 | 207110 | PF3D7_1405900 | RNA-binding protein, putative | Non_Synonymous_coding (GAT->CAT D518H) |
| PF3D7_14_v3 | 2029231 | | | Noncoding - upstream |

**Table S5.** ***In vitro*** **safety pharmacology of MMV693183**

| Toxicity | Test | Results |
|---|---|---|
| Off-target effects | Binding assay | All showed less than 25% inhibition/induction, except for GABA (non-selective) (25.2% induction) at 10 µM |
| | Enzyme and Uptake Assay | All showed less than 25% inhibition/induction, except for GABA transaminase (30.7% induction) and HDAC11 (36.9% inhibition) at 10 µM |
| | Patch clamp: hERG | $IC_{50}$ > 10 µM |
| | Patch clamp: $hNa_v$ 1.5 | $IC_{50}$ > 10 µM (Inhibition of 1.56% at 10 µM) |
| | Patch clamp: $hK_V$ 1.5 | $IC_{50}$ > 10 µM (Inhibition of 2.15% at 10 µM) |
| | Patch clamp: $hCa_V$ 1.2 | $IC_{50}$ > 10 µM (Inhibition of 24.20% at 10 µM) |
| Cytotoxicity | HepG2 viability | $IC_{50}$ > 10 µM |
| | Primary human and rat hepatocytes viability | $IC_{50}$ > 10 µM |
| Genotoxicity | AMES test | Non mutagenic in absence and presence of S-9 tested in 5 strains up to 5000 µg/plate. Differences in revertant numbers compared to control are depicted below<br><1.5-fold: TA102<br><2-fold: TA98, TA100<br><3-fold: TA1535, TA1537 |
| | Micronucleus Screening Assay | No induction of micronuclei in human peripheral blood lymphocytes up to 2000 µg/ml in presence or absence of S-9 (rat liver metabolizing system) |
| Phototoxicity | UV test | No absorbance >290 nm |
| Metabolism | CYP inhibition | $IC_{50}$ > 30 µM for human CYP1A2, CYP2D6, CYP3A4, CYP2C19 |
| | CYP induction | $IC_{50}$ > 10 µM for human CYP1A2, CYP2B6, CYP3A4 or rat CYP1A2, CYP2B1, CYP3A1 |

**Table S6. Study design seven-day repeat dose study in rats**

| Group No. | No. Animals | | | | Test material | Dose level (mg/kg/day) | Dose conc. (mg/ml) | Dose Volume (ml/kg) |
|---|---|---|---|---|---|---|---|---|
| | Toxicity | | Toxicokinetics | | | | | |
| | Male | Female | Male | Female | | | | |
| 1 | 3 | 3 | 2 | 2 | Control | 0 (Vehicle) | - | 10 |
| 2 | 3 | 3 | 2 | 2 | MMV693183 | 60 | 6 | 10 |
| 3 | 3 | 3 | 2 | 2 | MMV693183 | 200 | 20 | 10 |
| 4 | 3 | 3 | 2 | 2 | MMV693183 | 600 | 60 | 10 |

**Table S7. Toxicokinetic parameters in rats in the seven-day repeat dose study**

| | 60 mg/kg | | 200 mg/kg | | 600 mg/kg | |
|---|---|---|---|---|---|---|
| **Parameters** | **M** | **F** | **M** | **F** | **M** | **F** |
| **Day 1** | N = 2 | N = 2 | N = 2 | N = 2 | N = 2 | N = 2 |
| $t_{last}$ (h) | 8 | 8 | 8 | 8-24$^{\$}$ | 8-24$^{\$}$ | 8 |
| $t_{max}$ (h) | 0.25 | 0.25 | 0.25 | 0.25 | 0.25 | 0.25 |
| $C_{max}$ (µM) | 35.5 | 73.1 | 234 | 219 | 501 | 496 |
| $C_{max}$/D (kg*µM/mg) | 0.592 | 1.22 | 1.17 | 1.10 | 0.834 | 0.827 |
| $AUC_{0-8}$ (h*µM) | 41.2 | 88.9 | 345 | 320 | 990 | 1080 |
| $AUC_{0-8}$/D (h*kg*µM/mg) | 0.686 | 1.48 | 1.73 | 1.60 | 1.65 | 1.81 |
| $AUC_{last}$ (h*µM) | 41.2 | 88.9 | 345 | 333 | 1060 | 1080 |
| $AUC_{last}$/D (h*kg*µM/mg) | 0.686 | 1.48 | 1.73 | 1.67 | 1.77 | 1.81 |
| $AUC_{\infty}$ (h*µM) | n/a | 90.6 | 356 | 344 | 1120 | n/a |
| $AUC_{\infty}$/D (h*kg*µM/mg) | n/a | 1.51 | 1.78 | 1.72 | 1.86 | n/a |
| $t_{1/2}$ (h) | n/a | 1.36 | 1.55 | 2.97 | 2.78 | n/a |
| **Day 7** | N = 1* | N = 2 | N = 2 | N = 2 | N = 2 | N = 2 |
| $t_{last}$ (h) | 4 | 8 | 8 | 8 | 8 | 8-24$^{\$}$ |
| $t_{max}$ (h) | 0.25 | 0.25 | 0.25 | 0.25 | 0.25 | 0.25 |
| $C_{max}$ (µM) | 15.6 | 59.5 | 33.3 | 53.6 | 124 | 168 |
| $C_{max}$/D (kg*µM/mg) | 0.260 | 0.991 | 0.166 | 0.268 | 0.207 | 0.280 |
| $AUC_{0-8}$ (h*µM) | n/a | 69.7 | 46.2 | 54.9 | 147 | 150 |
| $AUC_{0-8}$/D (h*kg*µM/mg) | n/a | 1.16 | 0.231 | 0.274 | 0.244 | 0.251 |
| $AUC_{last}$ (h*µM) | 11.4 | 69.7 | 46.2 | 54.9 | 147 | 168 |
| $AUC_{last}$/D (h*kg*µM/mg) | 0.189 | 1.16 | 0.231 | 0.274 | 0.244 | 0.280 |
| $t_{1/2}$ (h) | n/a | 1.56 | 2.02 | 1.99 | n/a | 5.32 |

Mean data are presented of 2 female and 2 male mice per treatment: 60 mg/kg, 200 mg/kg, 600 mg/kg. N: number of animals; $: range; /D: dose normalized to 1 mg/kg; * Second male was excluded since only 2 measurable plasma concentrations were noted. Several $t_{1/2}$ values were not or could not be reported since the time span taken for the calculation of the half-life was not twice the calculated value or no log linear regression was possible. Consequently, $AUC_{\infty}$ could not be reported.

**Table S8. Body weight gain in rats from the seven-day repeat dose study**

| | Body weight gain (%) in males | | | | Body weight gain (%) in females | | | |
|---|---|---|---|---|---|---|---|---|
| | **0 mg/kg** | **60 mg/kg** | **200 mg/kg** | **600 mg/kg** | **0 mg/kg** | **60 mg/kg** | **200 mg/kg** | **600 mg/kg** |
| Day 1 | 0 (0.0) | 0 (0.0) | 0 (0.0) | 0 (0.0) | 0 (0.0) | 0 (0.0) | 0 (0.0) | 0 (0.0) |
| Day 4 | 11 (1.9) | 10 (0.9) | 10 (0.6) | 13 (1.0) | 7 (2.2) | 5 (0.9) | 7 (2.3) | 7 (1.9) |
| Day 7 | 14 (1.9) | 13 (1.0) | 13 (2.7) | 17 (1.8) | 6 (2.9) | 4 (2.2) | 7 (1.2) | 7 (1.6) |

**Table S9. Hematology and clinical chemistry parameters in the seven-day repeat dose study**

| | Male | | | | Female | | | |
|---|---|---|---|---|---|---|---|---|
| | **0 mg/kg** | **60 mg/kg** | **200 mg/kg** | **600 mg/kg** | **0 mg/kg** | **60 mg/kg** | **200 mg/kg** | **600 mg/kg** |
| **Hematology** | | | | | | | | |
| WBC 10E9/l | 8.6 (2.4) | 8.9 (0.9) | 6.4 (2.5) | 8.7 (2.0) | 5.2 (0.7) | 4.6 (1.5) | 6.0 (1.5) | 5.9 (1.4) |
| Neutrophils 10E9/l | 0.6 (0.1) | 0.6 (0.1) | 0.5 (0.2) | 0.7 (0.4) | 0.6 (0.1) | 0.4 (0.2) | 0.5 (0.1) | 0.6 (0.1) |
| Lymphocytes 10E9/l | 7.8 (2.1) | 8.1 (0.9) | 5.8 (2.3) | 7.8 (1.6) | 4.4 (0.6) | 4.1 (1.6) | 5.3 (1.5) | 5.2 (1.5) |
| Monocytes 10E9/l | 0.2 (0.1) | 0.1 (0.0) | 0.1 (0.1) | 0.1 (0.0) | 0.1 (0.0) | 0.0 (0.1) | 0.1 (0.1) | 0.1 (0.0) |
| Eosinophils 10E9/l | 0.1 (0.1) | 0.1 (0.1) | 0.0 (0.1) | 0.1 (0.1) | 0.1 (0.0) | 0.1 (0.1) | 0.1 (0.1) | 0.1 (0.0) |
| Basophils 10E9/l | 0.0 (0.0) | 0.0 (0.0) | 0.0 (0.0) | 0.0 (0.0) | 0.0 (0.0) | 0.0 (0.0) | 0.0 (0.0) | 0.0 (0.0) |
| Red blood cells 10E12/l | 7.16 (0.24) | 7.08 (0.29) | 6.80 (0.07) | 7.01 (0.31) | 7.46 (0.49) | 7.11 (0.45) | 7.61 (0.78) | 7.38 (0.21) |
| Reticulocytes 10e9/l | 380.0 (6.4) | 372.1 (10.1) | 363.2 (11.9) | 402.5 (29.0) | 230.7 (50.3) | 198.3 (17.6) | 226.7 (17.5) | 202.7 (50.7) |
| RDW % | 11.7 (0.7) | 11.9 (0.6) | 12.4 (1.4) | 12.6 (0.3) | 11.5 (1.0) | 10.7 (0.7) | 11.6 (0.8) | 11.4 (0.4) |
| Hemoglobin mmol/l | 9.2 (0.1) | 8.8 (0.2) | 8.4 (0.2) | 8.6 (0.5) | 8.8 (0.3) | 8.8 (0.8) | 9.1 (0.6) | 8.9 (0.3) |
| Hematocrit l/l | 0.444 (0.003) | 0.431 (0.009) | 0.404 (0.006) | 0.428 (0.027) | 0.426 (0.023) | 0.413 (0.033) | 0.432 (0.035) | 0.422 (0.004) |
| MCV fL | 62.0 (2.4) | 60.8 (1.4) | 59.5 (0.4) | 61.0 (1.1) | 57.2 (0.8) | 58.1 (1.4) | 56.8 (1.6) | 57.2 (1.0) |
| MCH fmol | 1.29 (0.02) | 1.24 (0.03) | 1.23 (0.03) | 1.23 (0.02) | 1.19 (0.04) | 1.23 (0.04) | 1.19 (0.06) | 1.20 (0.00) |
| MCHC mmol/l | 20.68 (0.44) | 20.44 (0.19) | 20.71 (0.23) | 20.13 (0.07) | 20.71 (0.44) | 21.12 (0.16) | 20.98 (0.43) | 21.03 (0.37) |
| Platelets 10e9/l | 907 (118) | 996 (29) | 1345 (77) | 1156 (190) | 937 (43) | 935 (145) | 1095 (184) | 1247 (33) |
| PT s | 20.2 (3.2) | 17.8 (1.5) | 18.0 (1.1) | 17.6 (1.3) | 18.9 (1.1) | 18.9 (0.6) | 18.1 (0.5) | 17.8 (0.8) |
| APTT s | 18.9 (0.2) | 16.1 (2.1) | 17.5 (3.6) | 17.9 (3.0) | 18.9 (3.9) | 16.9 (2.6) | 13.9 (1.3) | 15.1 (1.8) |
| **Clinical Chemistry** | | | | | | | | |
| ALAT U/l | 45.7 (8.9) | 48.9 (8.4) | 63.7 (31.0) | 54.2 (4.0) | 46.1 (17.3) | 37.4 (8.3) | 65.1 (7.1) | 64.9 (7.3) |
| ASAT U/l | 87.8 (9.4) | 84.5 (5.7) | 82.0 (7.5) | 76.1 (8.0) | 85.2 (5.2) | 97.6 (19.0) | 84.2 (3.4) | 85.0 (12.7) |
| ALP U/l | 336 (120) | 274 (33) | 394 (172) | 205 (39) | 222 (48) | 177 (29) | 151 (34) | 203 (68) |
| Total protein g/l | 58.9 (3.2) | 60.7 (1.7) | 62.9 (0.9) | 65.7 (2.0) | 62.1 (1.9) | 60.4 (2.4) | 62.7 (2.1) | 61.8 (1.7) |
| Albumin g/l | 31.1 (1.2) | 31.3 (0.2) | 32.2 (0.5) | 32.8 (0.5) | 34.3 (0.9) | 32.7 (1.7) | 32.5 (0.7) | 30.6 (1.3) |

| | | | | | | | | |
|---|---|---|---|---|---|---|---|---|
| Total globulin g/l | 27.7 (2.1) | 29.3 (1.5) | 30.8 (1.2) | 32.9 (1.5) | 27.8 (1.4) | 27.7 (0.7) | 30.2 (1.6) | 31.2 (0.5) |
| Alb/Gl ratio | 1.1 (0.0) | 1.1 (0.1) | 1.0 (0.1) | 1.0 (0.0) | 1.2 (0.1) | 1.2 (0.0) | 1.1 (0.1) | 1.0 (0.0) |
| Total bilirubin umol/l | 2.1 (0.4) | 1.4 (0.0) | 1.5 (0.2) | 1.6 (0.2) | 2.0 (0.1) | 1.9 (0.3) | 1.5 (0.1) | 1.5 (0.2) |
| Urea mmol/l | 8.4 (1.9) | 7.7 (1.3) | 8.6 (0.7) | 9.0 (0.6) | 7.5 (1.5) | 7.7 (0.9) | 8.1 (0.3) | 8.0 (1.6) |
| Creatinine umol/l | 32.4 (0.9) | 31.6 (2.5) | 32.4 (0.3) | 33.0 (1.3) | 33.0 (2.6) | 33.2 (1.1) | 33.3 (2.7) | 31.8 (0.7) |
| Glucose mmol/l | 8.39 (2.11) | 8.57 (1.31) | 9.11 (1.41) | 7.43 (1.64) | 7.35 (1.98) | 5.78 (0.41) | 6.03 (0.95) | 7.13 (0.63) |
| Cholesterol mmol/l | 2.33 (0.58) | 2.15 (0.18) | 2.02 (0.53) | 2.10 (0.25) | 1.67 (0.17) | 1.87 (0.31) | 2.85 (0.24) | 2.52 (0.18) |
| Triglycerides | 0.58 (0.16) | 0.72 (0.18) | 0.61 (0.09) | 0.58 (0.03) | 0.52 (0.09) | 0.38 (0.08) | 0.67 (0.21) | 0.49 (0.12) |
| Phospholipids mmol/l | 1.93 (0.58) | 1.78 (0.05) | 1.63 (0.36) | 1.85 (0.14) | 1.81 (0.18) | 1.72 (0.14) | 2.40 (0.04) | 2.15 (0.14) |
| Sodium mmol/l | 139.9 (1.3) | 140.8 (0.9) | 140.4 (0.4) | 139.7 (0.6) | 139.6 (0.7) | 140.2 (0.6) | 140.3 (1.3) | 139.3 (0.8) |
| Potassium mmol/l | 3.97 (0.34) | 3.99 (0.11) | 3.92 (0.03) | 4.19 (0.19) | 3.52 (0.33) | 3.76 (0.52) | 3.92 (0.16) | 3.82 (0.41) |
| Chloride mmol/l | 100 (1) | 102 (2) | 101 (1) | 101 (1) | 103 (2) | 105 (2) | 103 (1) | 103 (1) |
| Calcium mmol/l | 2.53 (0.05) | 2.56 (0.08) | 2.62 (0.02) | 2.65 (0.05) | 2.54 (0.04) | 2.54 (0.06) | 2.62 (0.11) | 2.57 (0.03) |
| Inorg.Phos mmol/l | 3.12 (0.11) | 2.71 (0.17) | 2.53 (0.12) | 2.78 (0.22) | 1.95 (0.37) | 2.32 (0.22) | 2.26 (0.27) | 2.20 (0.13) |

Mean and standard deviation values are presented for 3 male and 3 female mice per treatment. The haematology values from male and female control group and female 600 mg/kg group are determined in 2 mice only and therefore oneway ANOVA analysis was not performed on these values. Clinical Chemistry Parameters are measured in plasma. Cholesterol and phospholipids were increased, and albumin was decreased in female rats treated with higher doses, but the effects were only minor. Total protein, inorganic phosphate and total globulin remained within the normal range for rats of this age and strain.

**Table S10. Pharmacokinetic parameters in mice derived from data shown in Fig S12**

| **Dose** | **Route** | **Parameters** | **Units** | **Animal1** | **Animal2** | **Animal3** | **Mean** | **SD** |
|---|---|---|---|---|---|---|---|---|
| 3 | i.v. | $K_{el}$ | 1/hr | 0.59 | 0.55 | 0.61 | **0.58** | 0.03 |
| 3 | i.v. | $T_{1/2}$ | hr | 1.17 | 1.26 | 1.14 | **1.19** | 0.06 |
| 3 | i.v. | $C_0$ | ng/ml | 4944.5 | 5144.1 | 4952.3 | **5013.7** | 113.0 |
| 3 | i.v. | $AUC_{last}$ | hr*ng/ml | 3750.0 | 4121.9 | 3419.4 | **3763.8** | 351.5 |
| 3 | i.v. | $AUC_{0-\infty}$ | hr*ng/ml | 3752.1 | 4122.3 | 3419.8 | **3764.7** | 351.4 |
| 3 | i.v. | $CL_{total}$ | ml/min/kg | 13.33 | 12.13 | 14.62 | **13.36** | 1.25 |
| 3 | i.v. | Vss_obs | ml/kg | 1079.1 | 1028.1 | 972.3 | **1026.5** | 53.4 |
| 30 | p.o. | $K_{el}$ | 1/hr | 0.58 | 0.68 | 0.74 | **0.67** | 0.08 |
| 30 | p.o. | $T_{1/2}$ | hr | 1.20 | 1.02 | 0.93 | **1.05** | 0.14 |
| 30 | p.o. | $T_{max}$ | hr | 0.25 | 0.25 | 0.25 | **0.25** | 0.00 |
| 30 | p.o. | $C_{max}$ | ng/ml | 26100.0 | 25500.0 | 23700.0 | **25100.0** | 1249.0 |
| 30 | p.o. | $AUC_{last}$ | hr*ng/ml | 29452.7 | 27128.8 | 22505.2 | **26362.2** | 3536.6 |
| 30 | p.o. | $AUC_{0-\infty}$ | hr*ng/ml | 29453.4 | 27129.7 | 22505.7 | **26362.9** | 3536.7 |
| 30 | p.o. | $CL_{total}$ | ml/min/kg | 16.98 | 18.43 | 22.22 | **19.21** | 2.7 |
| 30 | p.o. | %F | % | 78.24 | 72.06 | 59.78 | **70.03** | 9.39 |

PK parameters were calculated using WinNonLin (Phoenix. Version 6.3). I.v.: intravenous; p.o.: per os.

**Table S11. Pharmacokinetic parameters in rats derived from data shown in Fig S12**

| **Dose** | **Route** | **Parameters** | **Units** | **Animal1** | **Animal2** | **Animal3** | **Mean** | **SD** |
|---|---|---|---|---|---|---|---|---|
| 3 | i.v. | $K_{el}$ | 1/hr | 0.19 | 0.23 | 0.25 | **0.22** | 0.03 |
| 3 | i.v. | $T_{1/2}$ | hr | 3.62 | 3.03 | 2.72 | **3.13** | 0.46 |
| 3 | i.v. | $C_0$ | ng/ml | 3682.2 | 2966.7 | 3086.4 | **3245.1** | 383.2 |
| 3 | i.v. | $AUC_{last}$ | hr*ng/ml | 2795.1 | 2252.0 | 2137.3 | **2394.8** | 351.4 |
| 3 | i.v. | $AUC_{0-\infty}$ | hr*ng/ml | 2802.5 | 2254.2 | 2138.5 | **2398.4** | 354.7 |
| 3 | i.v. | $CL_{total}$ | ml/min/kg | 17.84 | 22.18 | 23.38 | **21.14** | 2.91 |
| 3 | i.v. | Vss_obs | ml/kg | 1930.8 | 1963.6 | 1992.7 | **1962.4** | 31.0 |
| 30 | p.o. | $K_{el}$ | 1/hr | 0.27 | 0.28 | 0.26 | **0.27** | 0.01 |
| 30 | p.o. | $T_{1/2}$ | hr | 2.56 | 2.49 | 2.65 | **2.57** | 0.08 |
| 30 | p.o. | $T_{max}$ | hr | 0.25 | 0.25 | 0.25 | **0.25** | 0.00 |
| 30 | p.o. | $C_{max}$ | ng/ml | 21800.0 | 17200.0 | 22400.0 | **20466.7** | 2844.9 |
| 30 | p.o. | $AUC_{last}$ | hr*ng/ml | 33270.0 | 23500.9 | 30420.9 | **29063.9** | 5023.9 |
| 30 | p.o. | $AUC_{0-\infty}$ | hr*ng/ml | 33282.9 | 23509.2 | 30439.3 | **29077.1** | 5027.3 |
| 30 | p.o. | $CL_{total}$ | ml/min/kg | 15.02 | 21.27 | 16.43 | **17.57** | 3.28 |
| 30 | p.o. | %F | % | 138.77 | 98.02 | 126.92 | **121.24** | 20.96 |

PK parameters were calculated using WinNonLin (Phoenix. Version 6.3). I.v.: intravenous; p.o.: per os.

**Table S12. Pharmacokinetic parameters in dogs derived from data shown in Fig S12**

| Dose | Route | Parameters | Units | Animal1 | Animal2 | Animal3 | Mean | SD |
|---|---|---|---|---|---|---|---|---|
| 1 | i.v. | $T_{1/2}$ | hr | 2.15 | 5.69 | 4.83 | 4.23 | 1.85 |
| 1 | i.v. | $C_{max}$ | ng/ml | 1180 | 892 | 1160 | 1077 | 161 |
| 1 | i.v. | $AUC_{last}$ | hr*ng/ml | 1302 | 1514 | 1452 | 1423 | 109 |
| 1 | i.v. | $AUC_{0-\infty}$ | hr*ng/ml | 1347 | 1540 | 1466 | 1451 | 97.4 |
| 1 | i.v. | $CL_{total}$ | ml/min/kg | 12.37 | 10.82 | 11.37 | 11.52 | 0.787 |
| 1 | i.v. | Vss_obs | L/kg | 1.33 | 2.19 | 1.85 | 1.79 | 0.435 |
| 2 | p.o. | $T_{1/2}$ | hr | 2.11 | 4.97 | 1.70 | 2.93 | 1.78 |
| 2 | p.o. | $T_{max}$ | hr | 0.50 | 0.25 | 0.25 | 0.333 | 0.25-0.50 |
| 2 | p.o. | $C_{max}$ | ng/ml | 1630 | 1230 | 997 | 1286 | 320 |
| 2 | p.o. | $AUC_{last}$ | hr*ng/ml | 1942 | 1864 | 1727 | 1844 | 109 |
| 2 | p.o. | $AUC_{0-\infty}$ | hr*ng/ml | 2055 | 1880 | 1772 | 1902 | 143 |
| 2 | p.o. | F | % | 68.5 | 59.9 | 62.9 | 63.7 | 4.99 |

I.v.: intravenous; p.o.: per os.

**Table S13. Caco-2 permeability assay**

| Compound | Average Papp x10$^{-6}$/sec | | |
|---|---|---|---|
| | **A to B** | **B to A** | **Ratio** |
| MMV693183 | 2.9 | 13.5 | 4.7 |
| Furosemide | 0.2 | 12.9 | 67.1 |
| Atenolol | 0.4 | 0.4 | 1.0 |
| Verapamil | 19.0 | 40.3 | 2.1 |
| Carbamazepine | 25.0 | 28.9 | 1.2 |
| Domperidone | 3.5 | 25.2 | 7.3 |

Apical/Basal pH: 7.4/7.4

**Table S14.** ***In vitro*** **ADME parameters of MMV693183**

| | **Mouse** | **Rat** | **Dog** | **Human** |
|---|---|---|---|---|
| Fu,p | 0.666 | 0.601 | 0.587 | 0.481 |
| B:P | 0.98 | 0.91 | N.D. | 0.97 |
| *In vitro* $CL_{hep}$ (µl/min/$10^6$ cells) (SE) | 1.12 (2.02) | 3.35 (0.610) | 4.99 (1.17) | 0.40* |

*Data derived from Table 1

**Table S15. Metabolite identification in a human hepatocyte relay assay**

| Name | RT (min) | Ion (m/z) | Metabolism | *In vitro* (MS response) |
|---|---|---|---|---|
| Parent | 5.26 | 363 | - | 85% |
| M1 | 5.20 | 377 | Oxidation | 9% |
| M2 | 4.90 | 539 | Glucuronide | 2% |
| M3 | 4.69 | 383 (Na adduct) | Dehydrogenation | 2% |
| M4 | 5.03 | 383 (Na adduct) | Dehydrogenation | 2% |
| M5 | 4.69 | 539 | Glucuronide | <1% |
| M6 | 4.04 | 343 | Possibly loss of $H_2O$ | <1% |
| M7 | 4.21 | 443 | Sulphate | <1% |
| M8 | 4.59 | 539 | Glucuronide | <1% |
| M9 | 3.4.79 | 539 | Glucuronide | <1% |

RT: retention time

**Table S16. Metabolite identification in dog plasma and urine samples**

| Name | Ion (m/z) | RT (min) | Metabolism | Formula | Plasma | Urine |
|---|---|---|---|---|---|---|
| Parent | 362 | 3.36 | - | $C_{16}H_{21}F_3N_2O_4$ | Yes | Yes |
| M1 | M+174 | 2.89 | Glucuronide + dehydrogenation | $C_{22}H_{27}F_3N_2O_{10}$ | No | Yes |
| M2 | M+16 | 2.96 | Oxidation | $C_{16}H_{21}F_3N_2O_5$ | No | Yes |
| M3 | M+176 | 3.02 | Glucuronide | $C_{22}H_{29}F_3N_2O_{10}$ | No | Yes |
| M4 | M+174 | 3.07 | Glucuronide and Dehydrogenation | $C_{22}H_{27}F_3N_2O_{10}$ | No | Yes |
| M5 | M+176 | 3.14 | Glucuronide | $C_{22}H_{29}F_3N_2O_{10}$ | Yes | Yes |
| M6 | M-2 | 3.17 | Dehydrogenation | $C_{16}H_{19}F_3N_2O_4$ | Yes | Yes |
| M7 | M+172 | 3.22 | Glucuronide and didehydrogenation | $C_{22}H_{25}F_3N_2O_{10}$ | No | Yes |
| M8 | M-2 | 3.26 | Dehydrogenation | $C_{16}H_{19}F_3N_2O_4$ | Yes | Yes |
| M9 | M+14 | 3.36 | Oxidation and dehydrogenation | $C_{16}H_{19}F_3N_2O_5$ | Yes | Yes |
| M10 | M-4 | 3.65 | Tetra-dehydrogenation | $C_{16}H_{17}F_3N_2O_4$ | Yes | Yes |

**Table S17. Renal excretion of MMV693183 in rats intravenously treated with 3 mg/kg**

| Parameters | Units | Animal1 | Animal2 | Animal3 | Mean | SD |
|---|---|---|---|---|---|---|
| Stock concentration | mg/ml | 1.5 | 1.5 | 1.5 | | |
| Animal weight | g | 227 | 223 | 218 | | |
| Dose volume | µl | 460 | 440 | 440 | | |
| Dose administered | µg | 690 | 660 | 660 | | |
| Concentration in urine | ng/ml | 27920 | 106400 | 15600 | | |
| Urine volume | ml | 5.0 | 4.7 | 5.0 | | |
| Total urine excretion | µg | 139.6 | 500.1 | 78 | | |
| Urine fraction | | 0.20 | 0.76 | 0.12 | 0.36 | 0.35 |

**Table S18. Renal excretion of MMV693183 in dogs intravenously treated with 1 mg/kg**

| | Intervals (h) | Concentration (ng/ml) | Collected volume (ml) | Total in urine (µg) | Dose in urine |
|---|---|---|---|---|---|
| **Animal 1** | 0-4 | 2790 | 8 | 22.32 | |
| | 4-8 | - | 0 | - | |
| | 8-24 | - | 0 | - | |
| | Total | | 8 | 22.32 | 0.002 |
| **Animal 2** | 0-4 | 4550 | 60 | 273.0 | |
| | 4-8 | - | 0 | - | |
| | 8-24 | 6890 | 110 | 757.9 | |
| | Total | | 170 | 1030.9 | 0.089 |
| **Animal 3** | 0-4 | 7660 | 10 | 76.6 | |
| | 4-8 | - | 0 | - | |
| | 8-24 | 3700 | 6 | 22.2 | |
| | Total | | 16 | 98.8 | 0.009 |
| **Mean** | | | | 384 | 0.0332 |

In two dogs <20 ml of urine was collected and it is therefore likely that not all drug could be recovered in these animals. In the third dog, 170 ml of urine was collected and contains a higher fraction of compound (8.9%) than the other dogs and is therefore used for further studies.

**Table S19. Plasma clearance**

| Species | Plasma clearance (ml/min/kg) | | | | |
|---|---|---|---|---|---|
| | Predicted hepatic | Observed total | Observed renal | Observed total-renal = observed hepatic | Observed hepatic/ predicted hepatic |
| **Mouse** | 7.0 | 13.4 | n/d | <13.4 | <1.9 |
| **Rat** | 13.3 | 21 | 7.7 | 13.5 | 1.0 |
| **Dog** | 11.1 | 11.5 | 1.0 | 10.5 | 0.95 |
| **Human** | 0.51 | | | | |

Predicted hepatic clearance is based on the *in vitro* hepatic clearance data, Fu,p and B:P from table S14. The average of blood to plasma ratio in mouse, rat and human (0.95) was used in order to predict the hepatic clearance in dogs. The observed total clearance was derived from table S10-S12 and the observed renal clearance was determined from table S17-S18.

**Table S20. Human predicted PK parameters**

| Parameter | Value |
|---|---|
| CL, total (ml/min/kg) | 1.11 |
| CL, hepatic (ml/min/kg) | 0.51 |
| CL, renal (ml/min/kg) | 0.60 |
| V1 (l/kg) | 0.843 |
| V2 (l/kg) | 1.97 |
| Vss (l/kg) | 2.81 |
| Half-life (h) | 32.4 |

**Table S21. PK model estimates for the final PK model of MMV693183 from *in vivo* female NSG mice**

| PARAMETER | VALUE | RSE (%) | SHRINKAGE | COMMENT |
|---|---|---|---|---|
| Fabs0 | 1 (FIX) | - | - | Relative bioavailability (-) |
| CL | 1.25 | 41.70 | - | Apparent clearance (l/h) |
| Vc | 2.25 | 18.90 | - | Apparent central volume (l) |
| Q1 | 0.242 | 167 | - | Apparent intercompartmental clearance (l/h) |
| Vp1 | 799 | 347 | - | Apparent peripheral volume (l) |
| Q2 | 0.558 | 54.90 | - | Apparent intercompartmental clearance to second peripheral compartment (L/hour) |
| Vp2 | 1.81 | 28.70 | - | Apparent second peripheral volume (L) |
| Tk0 | 0.364 | 126 | - | Absorption time (h) |
| Tlag1 | 0 (FIX) | - | - | Absorption lag time (h) |
| **Inter-individual variability** | | | | |
| omega(Fabs0) | 0 (FIX) | - | - | Normal |
| omega(CL) | 0.354 | 18.90 | 3.30% | LogNormal |
| omega(Vc) | 0.195 | 56.70 | 57% | LogNormal |
| omega(Q1) | 0.3 (FIX) | - | 76% | LogNormal |
| omega(Vp1) | 0.3 (FIX) | - | 90% | LogNormal |
| omega(Q2) | 1.59 | 18.90 | 6.10% | LogNormal |
| omega(Vp2) | 0.343 | 36.50 | 38% | LogNormal |
| omega(Tk0) | 0.1 (FIX) | - | 94% | LogNormal |
| omega(Tlag1) | 0 (FIX) | - | - | Normal |
| **Residual Variability** | | | | |
| Additive error | 0.00143 | 32.50 | - | Compound concentration (ug/ml) |
| Proportional error | 0.235 | 12 | - | Fraction |
| Objective function | -265 | - | - | - |
| AIC | -239 | | | |
| BIC | -227 | | | |

**Table S22. Population model parameter estimates of the final PD model for MMV693183**

| PARAMETER | VALUE | RSE (%) | SHRINKAGE (%) | COMMENT |
|---|---|---|---|---|
| PLerr | 0 (FIX) | - | - | Individual deviation from baseline parasitemia |
| GR | 0.03 (FIX) | - | - | Net parasite growth rate (1/hour) |
| EMAX | 0.51 (FIX) | - | - | Maximum clearance rate (1/hour) |
| EC50 | 0.0115 | 37% | - | Concentration achieving 50percent of maximum effect (µg/mL) |
| hill | 2 (FIX) | - | - | Hill coefficient (.) |
| CLPara | 0.0698 | 6.75% | - | Parasitemia clearance (1/hour) |
| **Inter-individual variability** | | | | |
| omega(PLerr) | 0.2 (FIX) | - | 25% | Normal |
| omega(GR) | 0 (FIX) | - | - | LogNormal |
| omega(EMAX) | 0.1 (FIX) | - | 24% | LogNormal |
| omega(EC50) | 1.49 | 16.90% | 8.10% | LogNormal |
| omega(hill) | 0 (FIX) | - | - | Normal |
| omega(CLPara) | 0.0412 | 197% | 65% | LogNormal |
| **Parameter-Covariate relationships** | | | | |
| beta_CLPara(STUDYCOV_1) | -0.931 | 6.66% | - | Study covariate on CLPara |
| **Residual Variability** | | | | |
| error_ADD1 | 0.66 | 5.17 | - | Log-transformed parasitemia (percent or 1/ml) |
| Objective function | 1053 | - | - | - |
| AIC | 1061 | | | |
| BIC | 1065 | | | |

**Table 23. Human dose estimation based on the log total parasite reduction**

| **Parameter Criteria** | **Total Clearance prediction method** | **Median (mg)** | **AUC plasma (mg/l*h)** | **$C_{max}$ plasma (ng/ml)** |
|---|---|---|---|---|
| 9 log total parasite reduction | *in vitro* hepatocytes | 10 | 2.1 | 150 |
| 12 log total parasite reduction | *in vitro* hepatocytes | 15 | 3.2 | 230 |
| 9 log total parasite reduction | Allometry | 20 | 2.6 | 300 |
| 12 log total parasite reduction | Allometry | 30 | 3.9 | 450 |

**Table S24. Details on animal experiments**

| Study | Animals | Gender/ # animals | Age | Weight | Vehicle Control | Positive Control | Drug formulation | Doses (mg/kg/day) | Days of treatment | Route | Food |
|---|---|---|---|---|---|---|---|---|---|---|---|
| Single dose activity | NOD-SCID mice | F/8 | 6-8 weeks | 20-22 gram | No drug | 1x 50 mg/kg Chloroquine | Tween80-EtOH ((70/30-1:10 $H_2O$) | 50 | 1 | PO | *Ad libitum* |
| G6PD hemolysis | NOD-SCID mice | F/25* | 8 weeks | 19.6 gram average | PBS | 12.5 mg/kg Primaquine in PBS | Tween80-EtOH ((70/30-1:10 $H_2O$) | 10, 25, 50 | 4 | PO | *Ad libitum* |
| PKPD modeling | NOD-SCID mice | F/6 | 14-24 weeks | 25-28 gram | No drug | DHA | Tween80-EtOH ((70/30-1:10 $H_2O$) | 10, 25, 50, 100 | 1 | PO | *Ad libitum* |
| | | F/11 | 7 months | 22-28 gram | No drug | 2x10 mg/kg Chloroquine | Tween80-EtOH ((70/30-1:10 $H_2O$) | 1x2.5<br>1x5<br>2x2.5<br>2x5 | 3<br>1<br>3<br>3 | PO | *Ad libitum* |
| | | F/9 | 7 months | 22-28 gram | No drug | 2x10 mg/kg Chloroquine | Tween80-EtOH ((70/30-1:10 $H_2O$) | 5, 10, 25 | 1 | PO | *Ad libitum* |
| PKPD modeling | NOD-SCID mice | F/6 | 14-24 weeks | 25-28 gram | No drug | DHA | Tween80-EtOH ((70/30-1:10 $H_2O$) | 10, 25, 50, 100 | 1 | PO | *Ad libitum* |
| Seven-day repeat dose study in rats | Wistar Han rats | M/10, F/10 | 6 weeks | 126-194 gram | Elix water | N/A | Elix water | 60, 200, 600 | 7 | PO | *Ad libitum* |
| Pharmacokinetics mice | CD1 mice | M/6 | 6-8 weeks | 31-32 gram | N/A | N/A | 5% DMSO, 95% saline | 3 or 30 | 1 | IV or PO | *Ad libitum*# |

| | | | | | | | | | | | |
|---|---|---|---|---|---|---|---|---|---|---|---|
| Pharmacokinetics rats | Sprague Dawley rats | M/6 | | 250±50 gram | N/A | N/A | 5% DMSO, 95% saline for IV/5% DMSO, 95% saline, 0.5% carboxy methyl cellulose and 0.1% Tween-80 for PO | 3 or 30 | 1 | IV or PO | *Ad libitum*# |
| Pharmacokinetics dogs | Beagle dog | M/6 | 2-3 years | 8-11 kg | N/A | N/A | | 1 or 2 | 1 | IV or PO | |

* 1 mouse was excluded prior to engraftment injections due to small size and hunched appearance, 2 mice did not meet inclusion criteria (%HuRBCs > 60%).

#: Food fasting 12 h before dosing to 2 h after dosing. Gender F: female; M: male. Food fasting: fasting 12 h before dosing to 2 h after dosing; IV: intravenous; PO: *per os*.

**Table S25. Primers**

| Primer name | Primer sequence |
|---|---|
| **Sequencing & integration PCR primers** | |
| Sequencing guide | GCTTGGGGAGATCCGCC |
| Sequencing/Ct-Int-ACS-F | GGAAGAGATATAGATGGTACTGC |
| Sequencing-ACS-R | CGTCCAGATATCCAATAGTATCC |
| Ct-Int-ACS11-F | CAACCTATACGAAATAAATTTGGAGG |
| Ct-Int-R | CTCCAGTGAAAAGTTCTTCTCC |
| 3'-Int-F | TGTATCTATAGTTTCAAGTAGGACG |
| 3'-Int-ACS-R | AGGAGGGTATGATTTATTTCTTAAATC |
| 3'Int-ACS11-R | GGTACAAATATATGCATACGAGAAC |
| **ACS T648M point mutation primers** | |
| Guide-F | TATTGATTAATGCTCTTAAGGCTGT |
| Guide-R | aaacACAGCCTTAAGAGCATTAATc |
| Mutation-F | CAATTTTATATGGCTCCGACGGCATTAAGAG |
| Mutation-R | CTCTTAATGCCGTCGGAGCCATATAAAATTG |
| HR-F-EcoRI | tataaagaattcGGATCAACTGGTAAACCTAAAGG |
| HR-R-AatII | aatatagacgtcTCTGAACGCTCCATCTCC |
| **ACS-GFP primers** | |
| Ct-F-NotI | aattatgcggccgcCAGGGCACAGATTAGGAGC |
| Ct-R-MluI | aataatacgcgtTTTCTTAATTTCAATATGCTTTAACTTTTTTTTAC |
| 3'UTR-F-EcoRV-SacII | aattatgatatcccgcggTATATATATACCTTGTACTTACATTCGC |
| 3'UTR-R-BstZ17i | aatattgtatacGCGTGCTTTTTGATTTTTGCC |
| **ACS11-GFP primers** | |
| Ct-F-NotI | aataatgcggccgcTTTTCATCTCCCATATCAGAAGG |
| Ct-R-MluI | attattacgcgtCTTATTTCTTGATTTATAAAGATCATTTATTAC |
| 3'UTR-F-EcoRV-SacII | aatattgatatcccgcggAAAAGGGGTGAAATTAAAACAGACC |
| 3'UTR-R-BstZ17i | aataatgtatacGGCAAGAAAAATAAAGCAGAAATC |

Restriction sites are underlined
